# Evidence for the null hypothesis in functional magnetic resonance imaging using group-level Bayesian inference

**DOI:** 10.1101/2021.06.02.446711

**Authors:** Ruslan Masharipov, Yaroslav Nikolaev, Alexander Korotkov, Michael Didur, Denis Cherednichenko, Maxim Kireev

## Abstract

Classical null hypothesis significance testing is limited to the rejection of the point-null hypothesis; it does not allow the interpretation of non-significant results. Moreover, studies with a sufficiently large sample size will find statistically significant results even when the effect is negligible and may be considered practically equivalent to the ‘null effect’. This leads to a publication bias against the null hypothesis. There are two main approaches to assess ‘null effects’: shifting from the point-null to the interval-null hypothesis and considering the practical significance in the frequentist approach; using the Bayesian parameter inference based on posterior probabilities, or the Bayesian model inference based on Bayes factors. Herein, we discuss these statistical methods with particular focus on the application of the Bayesian parameter inference, as it is conceptually connected to both frequentist and Bayesian model inferences. Although Bayesian methods have been theoretically elaborated and implemented in commonly used neuroimaging software, they are not widely used for ‘null effect’ assessment. To demonstrate the advantages of using the Bayesian parameter inference, we compared it with classical null hypothesis significance testing for fMRI data group analysis. We also consider the problem of choosing a threshold for a practically significant effect and discuss possible applications of Bayesian parameter inference in fMRI studies. We argue that Bayesian inference, which directly provides evidence for both the null and alternative hypotheses, may be more intuitive and convenient for practical use than frequentist inference, which only provides evidence against the null hypothesis. Moreover, it may indicate that the obtained data are not sufficient to make a confident inference. Because interim analysis is easy to perform using Bayesian inference, one can evaluate the data as the sample size increases and decide to terminate the experiment if the obtained data are sufficient to make a confident inference. To facilitate the application of the Bayesian parameter inference to ‘null effect’ assessment, scripts with a simple GUI were developed.

## 1. Introduction

In the neuroimaging field, it is a common practice to identify statistically significant differences in local brain activity using the general linear model approach for mass-univariate null hypothesis significance testing (NHST) (Friston et al., 1994). NHST considers the probability of obtaining the observed data, or more extreme data, given that the null hypothesis of no difference is true. This probability, or p-value, of 0.01, means that, on average, in one out of 100 ‘hypothetical’ replications of the experiment, we find a difference no less than the one found under the null hypothesis. We conventionally suppose that this is unlikely, therefore, we ‘reject the null’; that is, NHST employs ‘proof by contradiction’ (Cohen, 1994). Conversely, when the p-value is large, it is tempting to ‘accept the null’. However, the absence of evidence is not evidence of absence (Altman and Bland, 1995). Using NHST, we can only state that we have ‘failed to reject the null’. Therefore, in the classical NHST framework, the question of interpreting non-significant results remains.

The most pervasive misinterpretation of non-significant results is that they provide evidence for the null hypothesis that there is no difference, or ‘no effect’ (Nickerson, 2000; Greenland et al., 2016; Wasserstein and Lazar, 2016). In fact, non-significant results can be obtained in two cases (Dienes, 2014): 1) the data are insufficient to distinguish the alternative from the null hypothesis, or 2) an effect is indeed null or trivial. To date, the extent to which the problem of making ‘no effect’ conclusions from non-significant results have affected the field of neuroimaging remains unclear, particularly in functional magnetic resonance imaging (fMRI) studies^1^. Regarding other fields of science such as psychology, neuropsychology, and biology, it was found that in 38–72% of surveyed articles, the null hypothesis was accepted based on non-significant results only (Finch et al., 2001; Schatz et al., 2005; Fidler et al., 2006; Hoekstra et al., 2006; Aczel et al., 2018).

Not mentioning non-significant results at all is another problem. Firstly, some authors may consider non-significant results disappointing or not worth publishing. Secondly, papers with non-significant results are less likely to be published. At the same time, NHST is usually based on the point-null hypothesis, that is, the hypothesis that the effect is *exactly* zero. However, the probability thereof is zero (Meehl, 1967; Friston et al., 2002a). This means that studies with a sufficiently large sample size will find statistically significant differences even when the effect is trivial or has no *practical* significance (Cohen, 1965, 1994; Serlin and Lapsley, 1985; Kirk, 1996). Therefore, ignoring non-significant results systematically biases our knowledge of true effects (Greenwald, 1975). This publishing bias is also known as the ‘file-drawer problem’ (Rosenthal, 1979; Ioannidis et al., 2014; De Winter and Dodou, 2015; for evidence in fMRI studies, see Jennings and Van Horn, 2012; Acar et al., 2018; David et al., 2018; Samartsidis et al., 2020).

Having the means to assess non-significant results would mitigate these problems. To this end, two main alternatives are available: Firstly, there are frequentist approaches that shift from point-null to interval-null hypothesis testing, for example, equivalence testing based on the two one-sided tests (TOST) procedure (Shuirmann, 1987; Wellek, 2010). Secondly, Bayesian approaches that are based on posterior parameter distributions (Lindley, 1965; Greenwald, 1975; Kruschke, 2010) and Bayes factors (Jeffreys, 1939/1948; Kass and Raftery, 1995; Rouder et al., 2009). The advantage of frequentist approaches is that they do not require a substantial paradigm shift (Campbell and Gustafson, 2018; Lakens, 2017). However, it has been argued that Bayesian approaches may be more natural and straightforward than frequentist approaches (Edwards et al., 1963; Lindley, 1975; Friston et al., 2002a; Wagenmakers, 2007; Rouder et al., 2009; Denies, 2014; Kruschke and Liddell, 2017b). It has long been noted that we tend to perceive lower p-values as stronger evidence for the alternative hypothesis, and higher p-values as evidence for the null, i.e., the ‘inverse probability’ fallacy as it is referred to by Cohen (1994). This is what we obtain in Bayesian approaches by calculating posterior probabilities. Instead of considering infinite ‘hypothetical’ replications and employing probabilistic ‘proof by contradiction’, Bayesian approaches directly provide evidence for the null and alternative hypotheses given the data, updating our prior beliefs in light of new relevant information. Bayesian inference allows us to ‘reject and accept’ the null hypothesis on an equal footing. Moreover, it allows us to talk about ‘low confidence’, indicating the need to either accumulate more data or revise the study design (see Fig. 1).

**Figure 1.**
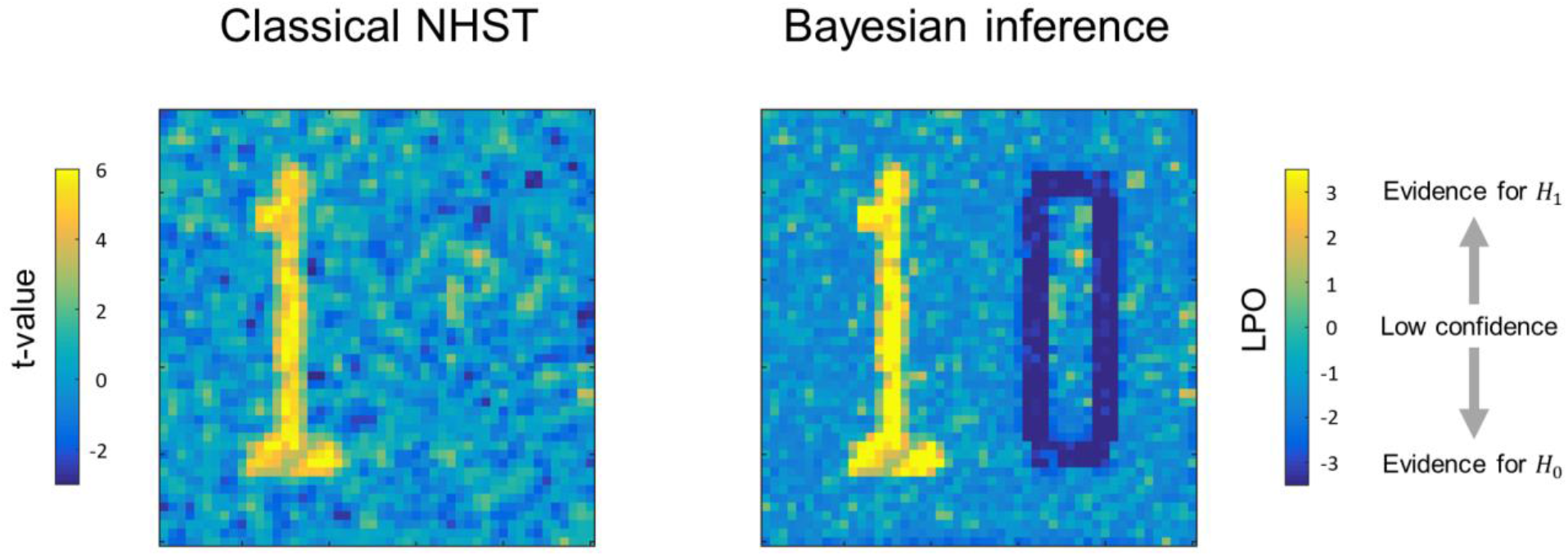
Possible results for the same data, obtained using classical NHST and Bayesian parameter inference. Classical NHST detects only areas with a statistically significant difference (‘number one’). Bayesian parameter inference based on the logarithm of posterior probability odds (*LPO*) provides us with additional information that is not available in classical NHST: a) it provides relative evidence for the null (*H*_0_) and alternative (*H*_1_) hypotheses, b) it detects areas with a trivial effect size (‘number zero’), c) it indicates ‘low confidence’ areas surrounding the ‘number one’ and ‘number zero’.

Despite the importance of this issue, and the high level of theoretical elaboration and implementation of Bayesian methods in common neuroimaging software programs, for example, Statistical Parametric Mapping 12 (SPM12) and FMRIB’s Software Library (FSL), to date, only a few fMRI studies implemented the Bayesian inference to assess ‘null effects’ (for example, see subject-level analysis in Magerkurth et al., 2015, group-level analysis in Dandolo and Schwabe, 2019; Feng et al., 2019). Therefore, this study is intended to introduce fMRI practitioners to the methods for assessing ‘null effects’. In particular, we focus on Bayesian parameter inference (Friston and Penny, 2003; Penny and Ridgway, 2013), as implemented in SPM12. Although Bayesian methods have been described elsewhere, the distinguishing feature of this study is that we aim to demonstrate the practical implementation of Bayesian inference to the assessment of ‘null effects’, and reemphasize its contributions over and above those of classical NHST. We deliberately aim to avoid mathematical details, which can be found elsewhere (Friston et al., 2002a, 2002b; Friston and Penny, 2003; Penny et al., 2003, 2005, 2007, 2013). Firstly, we briefly review the frequentist and Bayesian approaches for the assessment of the ‘null effect’. Next, we compare the classical NHST and Bayesian parameter inference on the Human Connectome Project (HCP) and the UCLA Consortium for Neuropsychiatric Phenomics datasets, focusing on group-level analysis. We then consider the choice of the threshold of the effect size for Bayesian parameter inference and estimate the typical effect sizes in different fMRI task designs. To demonstrate how the common sources of variability influence NHST and Bayesian parameter inference, we examine their behaviour for different sample sizes and spatial smoothing. Finally, we discuss practical research and clinical applications of Bayesian inference.

## 2. Theory

In this section, we briefly describe the classical NHST framework and review statistical methods which can be used to assess the ‘null effect’. We also considered two historical trends in statistical analysis: the shift from point-null hypothesis testing to interval estimation and interval-null hypothesis testing (Murphy and Myors, 2004; Wellek, 2010; Cumming, 2013), and the shift from frequentist to Bayesian approaches (Kruschke and Liddell, 2017b).

### 2.1. Classical NHST framework

Most task-based fMRI studies rely on the general linear model approach (Friston et al., 1994; Poline and Brett, 2012). It provides a simple way to separate blood-oxygenated-level dependent (BOLD) signals associated with particular task conditions from nuisance signals and residual noise when analysing single-subject data (subject-level analysis). At the same time, it allows us to analyse BOLD signals within one group of subjects or between different groups (group-level analysis). Firstly, we must specify a general linear model and estimate its parameters (*β* values). Some of these parameters reflect the mean amplitudes of BOLD responses evoked in particular task conditions or groups of subjects. The linear contrast of these parameters, *θ* = *cβ*, represents the experimental effect of interest (hereinafter ‘*the effect*’), expressed as the difference between two conditions or groups of subjects. Next, we test the effect against the point-null hypothesis, *H*_0_: *θ* = *γ* (usually, *θ* = 0). To do this, we use test statistics that summarise the data in a single value, for example, the t-value. For the one-sample case, the t-value is the ratio of the discrepancy of the estimated effect from the hypothetical null value to its standard error. Thus, it represents the contrast-to-noise ratio, which is similar to the signal-to-noise ratio (Welvaert and Rosseel, 2013). Finally, we calculate the probability of obtaining the observed t-value or a more extreme value, given that the null hypothesis is true (p-value). This is also commonly formulated as the probability of obtaining the observed data or more extreme data, given that the null hypothesis is true (Cohen, 1994). It can be simply written as a conditional probability *P*(*D* + *H*_0_), where ‘*D* +’ denotes the observed data or more extreme data which can be obtained in infinite ‘hypothetical’ replications under the null (Schneider, 2014, 2018). If this probability is lower than some conventional threshold, or alpha level (for example, *α* = 0.05), then we can ‘reject the null hypothesis’ and state that we found a statistically significant effect. When this procedure is repeated for a massive number of voxels, it is referred to as ‘mass-univariate analysis’. However, if we consider *m* = 100 000 voxels with no true effect and repeat significance testing for each voxel at α = 0.05, we would expect to obtain 5000 false rejections of the null hypothesis (false positives). To control the number of false positives, we must reduce the alpha level for each significance test by applying the multiple comparison correction (Worsley et al., 1992; Genovese et al., 2002; Nichols and Hayasaka, 2003; Nichols, 2012).

To date, the classical NHST has been the most widely used statistical inference method in neuroscience, psychology, and biomedicine (Szucs and Ioannidis, 2017, 2020; Ioannidis, 2019). It is often criticised for the use of the point-null hypothesis (Meehl, 1967), also known as the ‘nil null’ (Cohen, 1994) or ‘sharp null’ hypothesis (Edwards et al., 1963). It was argued that the point-null hypothesis could be appropriate only in hard sciences such as physics, but it is always false in soft sciences; this problem is sometimes known as the Meehl’s paradox (Meehl, 1967, 1978; Serlin and Lapsley, 1985, 1993; Cohen, 1994; Kirk, 1996). In the case of fMRI research, we face complex brain activity which is influenced by numerous psychophysiological factors. This means that with a large amount of data, we find a statistically significant effect in all voxels for any linear contrast (Friston, 2002a). For example, Gonzalez-Castillo et al. (2012) showed a statistically significant effect in over 95% of the brain for simple visual stimulation when averaging single-subject data from 100 runs (approximately four hours of scanning). Approximately half of the brain areas showed activation, whereas the other half showed deactivation. Whole-brain (de)activations can also be found when analysing large datasets such as the HCP (*N* ~ 1000) or UK Biobank (*N* ~ 10 000) datasets. When we increase the sample size, the effect estimate does not change much. Still, the standard error in the denominator of the t-value becomes increasingly smaller, resulting in negligible effects becoming statistically significant. Thus, the classical NHST ignores the magnitude of the effect. Attempts to overcome this problem led to the proposal of making a distinction between ‘statistical significance’ and ‘material significance’ (Hodges and Lehmann, 1954) or ‘practical significance’ (Cohen, 1965; Kirk, 1996). That is, we can test whether the effect size is larger or smaller than some practically meaningful value using interval-null hypothesis testing (Friston et al., 2002a, 2002b, 2013).

### 2.2. Frequentist approach to interval-null hypothesis testing

Interval-null hypothesis testing is widely used in medicine and biology (Meyners, 2012). Consider, for example, a pharmacological study designed to compare a new treatment with an old treatment that has already shown its effectiveness. Let *β_new_* be the mean effect on brain activity of the new treatment and *β_old_* the mean effect of the old treatment. Then, *θ* = (*β_new_* – *β_old_*) is the relative effect of the new treatment. The practical significance is defined by the effect size (ES) threshold *γ*. If a larger effect on brain activity is preferable, then we can test whether there is a practically meaningful difference in a positive direction (*H*_1_: *θ* > *γ* vs. *H*_0_: *θ* ≤ *γ*). This procedure is known as the *superiority test* (see Fig. 2A). We can also test whether the effect of the new treatment is no worse (practically smaller) than the effect of the old treatment (*H*_1_: *θ* > −*γ* vs. *H*_0_: *θ* ≤ −*γ*). This procedure is sometimes known as the *non-inferiority test* (see Fig. 2B). If a smaller effect on brain activity is preferable, we can use the superiority or non-inferiority test in the opposite direction (see Fig. 2C–D). The combination of these two superiority tests allows us to find a practically meaningful difference in both directions (*H*_1_: *θ* > *γ* and *θ* < −*γ* vs. *H*_0_: −*γ* ≤ *θ* ≤ *γ*), that is, the *minimum-effect test* (see Fig. 2E). The combination of the two non-inferiority tests allows us to reject the hypothesis of practically meaningful differences in any direction (*H*_1_: −*γ* ≤ *θ* ≤ *γ* vs. *H*_0_: *θ* > *γ* and *θ* < −*γ*). This is the most widely used approach to *equivalence testing*, known as the *two one-sided tests* (TOST) procedure (see Fig. 2F). For more details on the superiority and minimum-effect tests, see (Serlin and Lapsey, 1985, 1993; Murphy and Myors, 1999, 2004). For more details on the non-inferiority test and TOST procedure see (Schuirmann, 1987; Rogers et al., 1993; Wellek, 2010; Meyners, 2012; Lakens, 2017).

**Figure 2.**
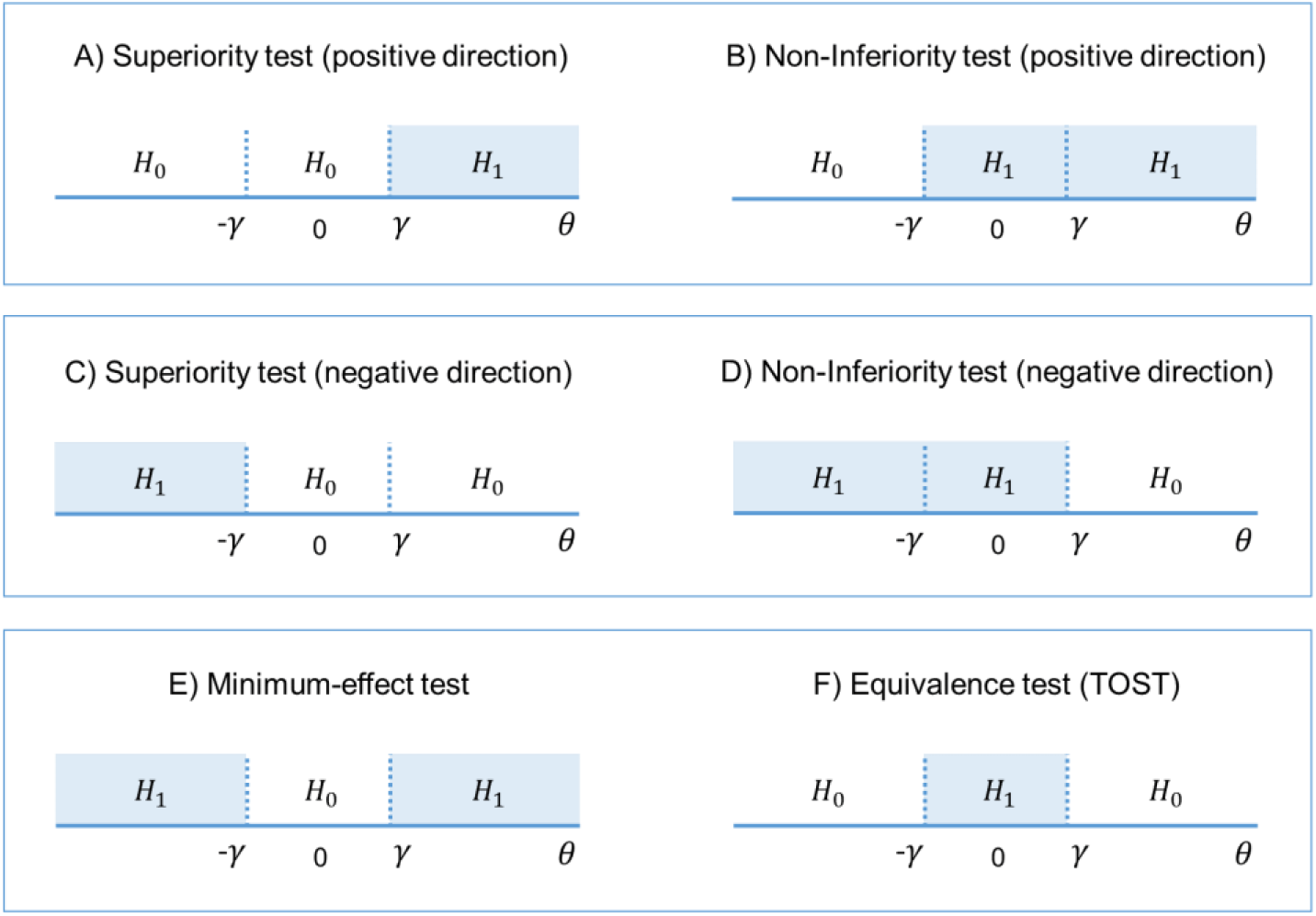
The alternative (*H*_1_) and null (*H*_0_) hypotheses for different types of interval-null hypotheses tests. A–B) One-sided tests in the positive direction (‘the larger is better’). C–D) One-sided tests in the negative direction (‘the smaller is better’). E) Combination of both superiority tests. F) Combination of both non-inferiority tests. Scheme modified from Aisbett et al. (2020)

The interval [−*γ; γ*] defines trivially small effect sizes that we consider to be equivalent to the ‘null effect’ for practical purposes. This interval is also known as the ‘equivalence interval’ (Schuirmann, 1987) or ‘region of practical equivalence (ROPE)’ (Kruschke, 2011). The TOST procedure, in contrast to classical NHST, allows us to assess the ‘null effects’. If we reject the null hypothesis of a practically meaningful difference, we can conclude that the effect is trivially small. The TOST procedure can also be intuitively related to frequentist interval estimates, known as confidence intervals (‘confidence interval approach’, Westlake, 1972; Schuirmann, 1987). Confidence intervals reflect the uncertainty in the point estimation of the parameters defined by its standard error. The confidence level of (1 – *α*) means that among infinite ‘hypothetical’ replications, (1 – *α*)% of the confidence intervals will contain the true effect under the null. Therefore, the TOST procedure is operationally identical to considering whether the (1 – 2*α*)% confidence interval falls entirely into the ROPE, as it uses two one-sided tests with an alpha level of *α*.

Interval-null hypothesis testing can be used in fMRI studies not only to compare the effects of different treatments. For example, we can apply superiority tests in the positive and negative directions to detect ‘activated’ and ‘deactivated’ voxels and additionally apply the TOST procedure to detect ‘not activated’ voxels. However, even though we can solve the Meehl’s paradox and assess the ‘null effects’ by switching from point-null to interval-null hypothesis testing within the frequentist approach, this approach still has fundamental philosophical and practical difficulties which can be effectively addressed using Bayesian statistics.

### 2.3. Pitfalls of the frequentist approach

The pitfalls of the frequentist approach have been actively discussed by statisticians and researchers for decades. We believe that the Bayesian approach is clearer and more coherent (Lindley, 1990). Unlike the frequentist approach, it does not require special modifications or adjustments to achieve similar practical goals. Here, we briefly mention a few of the main problems associated with the frequency approach.

1. NHST is a hybrid of Fisher’s approach that focuses on the p-value (thought to be a measure of evidence against the null hypothesis), and Neyman-Pearson’s approach that focuses on controlling false positives with the alpha level while maximising true positives in long-run replications. These two approaches are argued to be incompatible and have given rise to several misinterpretations among researchers, for example, confusing the meaning of p-values and alpha levels (Edwards et al., 1963; Gigerenzer, 1993; Goodman, 1993; Finch et al., 2001; Berger, 2003; Hubbard and Bayarri, 2003; Royall, 1997; Turkheimer et al., 2004, Perezgonzalez, 2015; Schneider, 2014; Szucs and Ioannidis, 2017; Greenland, 2019).
2. The logical structure of NHST is the same as that of ‘proof by contradiction’ or ‘indirect proof’, which becomes formally invalid when applied to probabilistic statements (Pollard and Richardson, 1987; Cohen, 1994; Falk and Greenbaum, 1995; Nickerson, 2000; Sober, 2008; Schneider, 2014, 2018; Wagenmakers et al., 2017; but see Hagen, 1997). Valid ‘proof by contradiction’ can be expressed in syllogistic form as: 1) ‘If A, then B’ (Premise N<u>o</u>1), 2) ‘Not B’ (Premise N<u>o</u>2), 3) ‘Therefore not A’ (Conclusion). Probabilistic ‘proof by contradiction’ in relation to NHST can be formulated as: 1) ‘If *H*_0_ is true, then *D* + are highly unlikely, 2) ‘*D* + was obtained’, 3) ‘Therefore *H*_0_ is highly unlikely’. This problem is also referred to as the ‘illusion of probabilistic proof by contradiction’ (Falk and Greenbaum, 1995). To illustrate the fallacy of such logic, consider the following example from Pollard and Richardson (1987): 1) ‘If a person is an American (*H*_0_), then he is most probably not a member of Congress’, 2) ‘The person is a member of Congress’, 3) ‘Therefore the person is most probably not an American’. Based on this, one ‘rejects the null’ and makes an obviously wrong inference, as only American citizens can be a member of Congress. At the same time, using Bayesian statistics, we can show that the null hypothesis (‘the person is an American’) is true (see the Bayesian solution of the ‘Congress example’ in the Supplementary Materials). The ‘illusion of probabilistic proof by contradiction’ leads to widespread confusion between the probability of obtaining the data, or more extreme data, under the null *P*(*D* + |*H*_0_) and the probability of the null under the data *P*(*H*_0_|D) (Pollard and Richardson, 1987; Gigerenzer, 1993; Cohen, 1994; Falk and Greenbaum, 1995; Nickerson, 2000; Finch et al., 2001; Hoekstra et al., 2006; Goodman, 2008; Wasserstein and Lazar, 2016; Greenland, 2016; Amrhein and Roth, 2017). The latter is a posterior probability calculated based on Bayes’ rule. The fact that researchers usually treat the p-value as a continuous measure of evidence (the Fisherian interpretation) only exacerbates this problem. ‘The lower the p-value, the stronger the evidence against the null’ statement can be erroneously transformed to statements such as ‘the lower the p-value, the stronger the evidence for the alternative’ or ‘the higher the p-value, the stronger the evidence for the null’. NHST can only provide evidence *against*, but never *for*, a hypothesis. In contrast, posterior probability provides direct evidence for a hypothesis; hence, it has a simple intuitive interpretation.
3. The p-value is not a plausible measure of evidence (Cornfield, 1966; Royall, 1986, 1997; Berger and Sellke, 1987; Berger and Berry, 1988; Goodman, 1993; Wagenmakers et al., 2007, 2008, 2017; Hubbard and Lindsay, 2008; Johansson, 2011; Wasserstein and Lazar, 2016; bet see Greenland, 2019). The frequentist approach considers infinite ‘hypothetical’ replications of the experiment (sampling distribution); that is, the p-value depends on unobserved data. One of the most prominent theorists of Bayesian statistics, Harold Jeffreys, put it as follows: ‘*What the use of P implies, therefore, is that a hypothesis that may be true may be rejected because it has not predicted observable results that have not occurred”* (Jeffreys, 1948, p. 357). In turn, the sampling distribution depends on the researcher’s intentions. These intentions may include different kinds of *multiplicities*, such as multiple comparisons, double-sided comparisons, secondary analyses, subgroup analyses, exploratory analyses, preliminary analyses, and interim analyses of sequentially obtained data with optional stopping (Gopalan and Berry, 1998). Two researchers with different intentions may obtain different p-values based on the same dataset. The problem is that these intentions are usually unknown. When null findings are considered disappointing, it is tempting to increase the sample size until one obtains a statistically significant result. However, a statistically significant result may arise when the null is, in fact, true, which can be shown by Bayesian statistics. That is, the p-value usually exaggerates evidence against the null hypothesis. The discrepancy that may arise between frequentist and Bayesian inference is also known as the Jeffreys–Lindley paradox (Jeffreys, 1939/1948; Lindley, 1957). In addition, it is argued that a consistent measure of evidence should not depend on the sample size (Cornfield, 1966). However, identical p-values provide different evidence against the null hypothesis for small and large sample sizes (Wagenmakers, 2007). In contrast, evidence provided by posterior probabilities and Bayes factors does not depend on the testing or stopping intentions or the sample size (Kruschke and Liddell, 2017b; Wagenmakers, 2007).
4. Although frequentist interval estimates (Cohen, 1990, 1994; Cumming, 2013) and interval-based hypothesis testing (Murphy and Myors, 2004; Wellek, 2010; Meyners, 2012; Lakens, 2017) greatly facilitate the mitigation of the abovementioned pitfalls in data interpretation, they are still subject to some of the same types of problems as the p-values and classic NHST (Cortina and Dunlap, 1997; Nickerson, 2000; Belia et al., 2005; Wagenmakers et al., 2008; Kruschke, 2013; Kruschke and Liddell, 2017a; Hoekstra et al., 2014; Morey et al., 2015; Greenland et al., 2016). Confidence intervals also depend on unobserved data and the intentions of the researcher. Moreover, the meaning of confidence intervals seems counterintuitive to many researchers. For example, one of the most common misinterpretations of the (1 – *α*)% confidence interval is that the probability of finding an effect within the confidence interval is (1 – *α*)%. In fact, it is a Bayesian interval estimate known as a *credible* interval.

Nevertheless, we would like to emphasise that we do not advocate abandoning the frequency approach. Correctly interpreted frequentist interval-based hypothesis testing with a priori power analysis defining the sample size and proper multiplicity adjustments often lead to conclusions similar to those of Bayesian inference (Lakens et al., 2018). However, even honest researchers with transparent intentions may find it logically and practically difficult to carry out an appropriate power analysis and make multiplicity adjustments (Berry and Hochberg, 1999; Cramer et al., 2015; Schönbrodt et al., 2015; Streiner, 2015; Sjölander and Vansteelandt, 2019). These procedures may be even more complicated in fMRI research than in psychological or social studies (see discussion on power analysis in Mumford and Nichols, 2008; Joyce and Hayasaka, 2012; Mumford, 2012; Cremes et al., 2017; Poldrack et al., 2017; multiple comparisons in Nichols and Hayasaka, 2003; Nichols, 2012; Eklund et al., 2016; and other types of multiplicities in Turkheimer et al., 2004; Chen et al., 2018, 2019, 2020; Alberton et al., 2020). For example, at the beginning of a long-term study, one may want to check whether stimulus onset timings are precisely synchronised with fMRI data collection and perform preliminary analysis on the first five subjects. The question of whether the researcher must make an adjustment for this technical check when reporting the results for the final sample become important in the frequentist approach. Such preliminary analyses (or other forms of interim analyses) are not a source of concern in Bayesian inference. Or, for example, one may want to find both ‘(de)activated’ and ‘not activated’ brain areas and use two superiority tests in combination with the TOST procedure. It is not trivial to make appropriate multiplicity adjustments in this case. In contrast, Bayesian inference suggests a single decision rule without the need for additional adjustments. Moreover, to our knowledge, practical implementations of superiority tests and the TOST procedure in common software for fMRI data analysis do not yet exist. At the same time, Bayesian analysis has already been implemented in SPM12 (https://www.fil.ion.ucl.ac.uk/spm/software/spm12) and is easily accessible to end-users. It consists of two steps: Bayesian parameter estimation and Bayesian inference. In general, it is not necessary to use Bayesian analysis at the subject level of analysis to apply it at the group level. One can combine computationally less demanding frequentist parameter estimation for single subjects with Bayesian estimation and inference at the group level. In the next sections, we consider the group-level Bayesian analysis implemented in SPM12.

### 2.4. Bayesian parameter estimation

Bayesian statistics is based on Bayes’ rule:

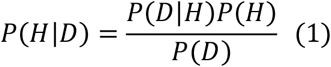

where *P*(*H*|*D*) is the probability of the hypothesis given the obtained data or posterior probability. *P*(*D*|*H*) is the probability of obtaining the *exact* data given the hypothesis or the likelihood (notice the difference from *P*(*D* + |*H*), which includes *more extreme* data). *P*(*H*) is the prior probability of the hypothesis (our knowledge of the hypothesis before we obtain the data). *P*(*D*) is a normalising constant ensuring that the sum of posterior probabilities over all possible hypotheses equals one. For example, if we consider two mutually exclusive hypotheses *H*_0_ and *H*_1_, then *P*(*D*) = *P*(*D*|*H*_0_) *P*(*H*_0_) + *P*(*D*|*H*_1_)*P*(*H*_1_) and *P*(*H*_0_|*D*) + *P*(*H*_1_|*D*) = 1.

In verbal form, Bayes’ rule can be expressed as:

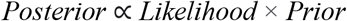

This means that we can update our prior beliefs about the hypothesis based on the obtained data. The main difficulty in using Bayesian statistics lies in choosing the appropriate prior assumptions. If the data are organised hierarchically, which is the case for neuroimaging data, priors can be specified based on the data using an empirical Bayesian approach. The lower level of the hierarchy corresponds to the experimental effects at any given voxel, and the higher level of the hierarchy comprises the effect over all voxels. Thus, the variance of the experimental effect over all voxels can be used as the prior variance of the effect at any given voxel. This approach is known as the parametric empirical Bayes (PEB) with the ‘global shrinkage’ prior (Friston and Penny, 2003). The prior variance is estimated from the data under the assumption that the prior probability density corresponds to a Gaussian distribution with zero mean. In other words, a global experimental effect is assumed to be absent. An increase in local activity can be detected in some brain areas; a decrease can be found in others, but the total change in neural metabolism in the whole brain is approximately zero. This is a reasonable physiological assumption because studies of brain energy metabolism have shown that the global metabolism is ‘remarkably constant despite widely varying mental and motoric activity’ (Raichle and Gusnard, 2002), and ‘the changes in the global measurements of blood flow and metabolism’ are ‘too small to be measured’ by functional imaging techniques such as PET and fMRI (Gusnard and Raichle, 2001).

Now, we can rewrite Bayes’ rule (eq. 1) for the effect *θ* = *cβ*:

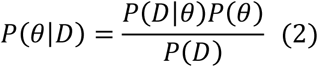

In the process of Bayesian updating with the ‘global shrinkage’ prior, the effect estimate ‘shrinks’ toward zero. The greater the uncertainty of the effect estimate (variability) in a particular voxel, the less confidence in this estimate, and the more it shrinks (see Fig. 3).

**Figure 3.**
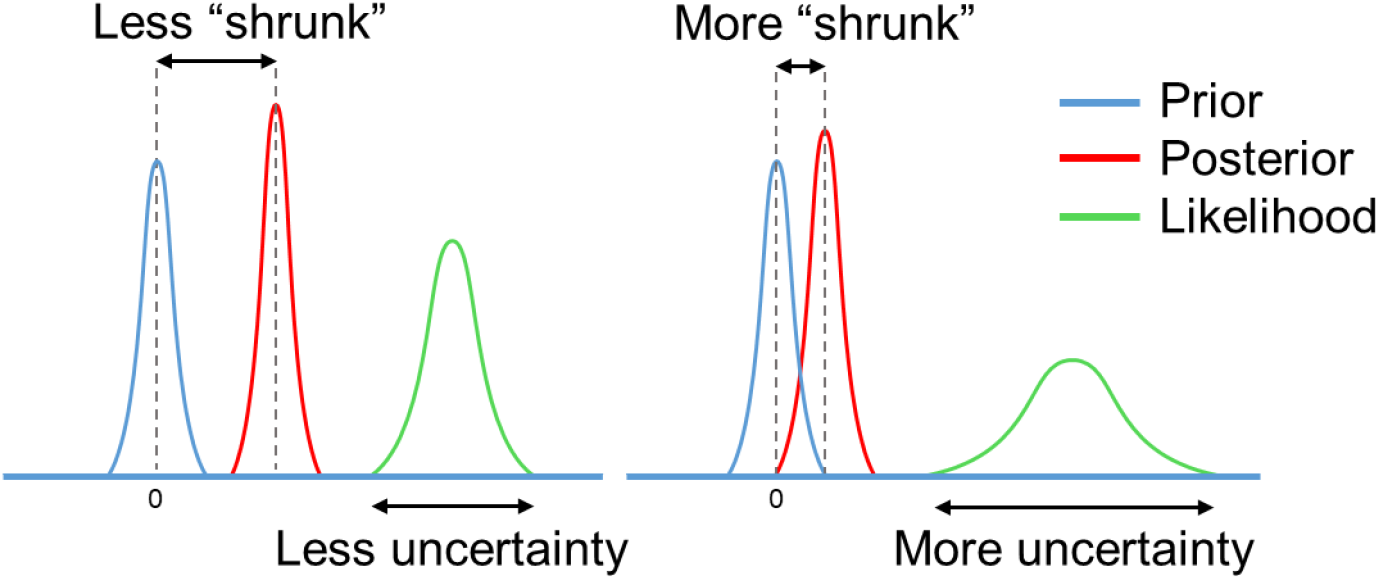
Schematic of Bayesian updating with the ‘global shrinkage’ prior. Scheme modified from Stephan (2016).

After Bayesian parameter estimation, we can apply one of the two main types of Bayesian inference (Penny and Ridgway, 2013): *Bayesian parameter inference (BPI)* or *Bayesian model inference (BMI)*. BPI is also known as Bayesian parameter estimation (Kruschke and Liddell, 2017b). However, we deliberately separate these two terms, as they correspond to two different steps of data analysis in SPM12. BMI is also known as Bayesian model comparison, Bayesian model selection, or Bayesian hypothesis testing (Kruschke and Liddell, 2017b). We chose the term BMI as it is consonant with the term BPI.

### 2.5. Bayesian parameter inference

The BPI is based on the posterior probability of finding the effect within or outside the ROPE. Let effects larger than the ES threshold *γ* be ‘activations’, those smaller than –*γ* be ‘deactivations’, and those falling within the ROPE [−*γ*; *γ*] be ‘no activations’. Then, we can classify voxels as ‘activated’, ‘deactivated’, or ‘not activated’ if:

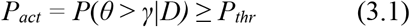

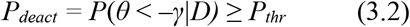

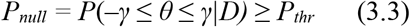

where *P_thr_* is the posterior probability threshold (usually *P_thr_* = 95%). Note that *P_act_* + *P_deact_* + *P_null_* 1.

If none of the above criteria are satisfied, the data in a particular voxel are insufficient to distinguish voxels that are ‘(de)activated’ from those that are ‘not activated’. Hereinafter, we refer to them as ‘low confidence’ voxels (Magerkurth et al., 2015). This decision rule is also known as the ‘ROPE-only’ rule (Kruschke and Liddell, 2017a, see also Greenwald (1975), Wellek (2010), Liao et al. (2019). To the best of our knowledge, the application of this decision rule to neuroimaging data was pioneered by Friston et al. (2002a, 2002b, 2003). For convenience and visualisation purposes, we can use the natural logarithm of the posterior probability odds (LPO), for example:

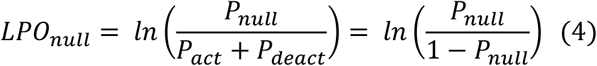

This allows us to more effectively discriminate voxels with a posterior probability close to unity (Penny and Ridgway, 2013). *LPO_null_* > 3 corresponds to *P_null_* > 95%. In addition, *LPO* also allows us to identify the connection between BPI and BMI. The maps of the *LPO* are termed posterior probability maps (PPMs) in SPM12.

Another possible decision rule considers the overlap between ROPE and the 95% highest density interval (HDI). HDI is a type of credible interval (Bayesian analogue of the confidence interval), which contains only the effects with the highest posterior probability density. If the HDI falls entirely inside the ROPE, we can classify voxels as ‘not activated’. In contrast, if the HDI lies completely outside the ROPE, we can classify voxels as either ‘activated’ or ‘deactivated’. If the HDI overlaps with the ROPE, we cannot make a confident decision (we can consider them to be ‘low confidence’ voxels). This decision rule is known as the ‘HDI+ROPE’ rule (Kruschke and Liddell, 2017a). It is more conservative than the ‘ROPE-only’ rule, as it does not consider the effects from the tails of a posterior probability distribution. When the posterior distributions are skewed, the ‘HDI+ROPE’ decision rule should be used. However, in SPM12, the ‘HDI+ROPE’ and ‘ROPE-only’ decision rules should produce similar results because SPM12 utilises normal distributions. These decision rules are illustrated in Fig. 4.

**Figure 4.**
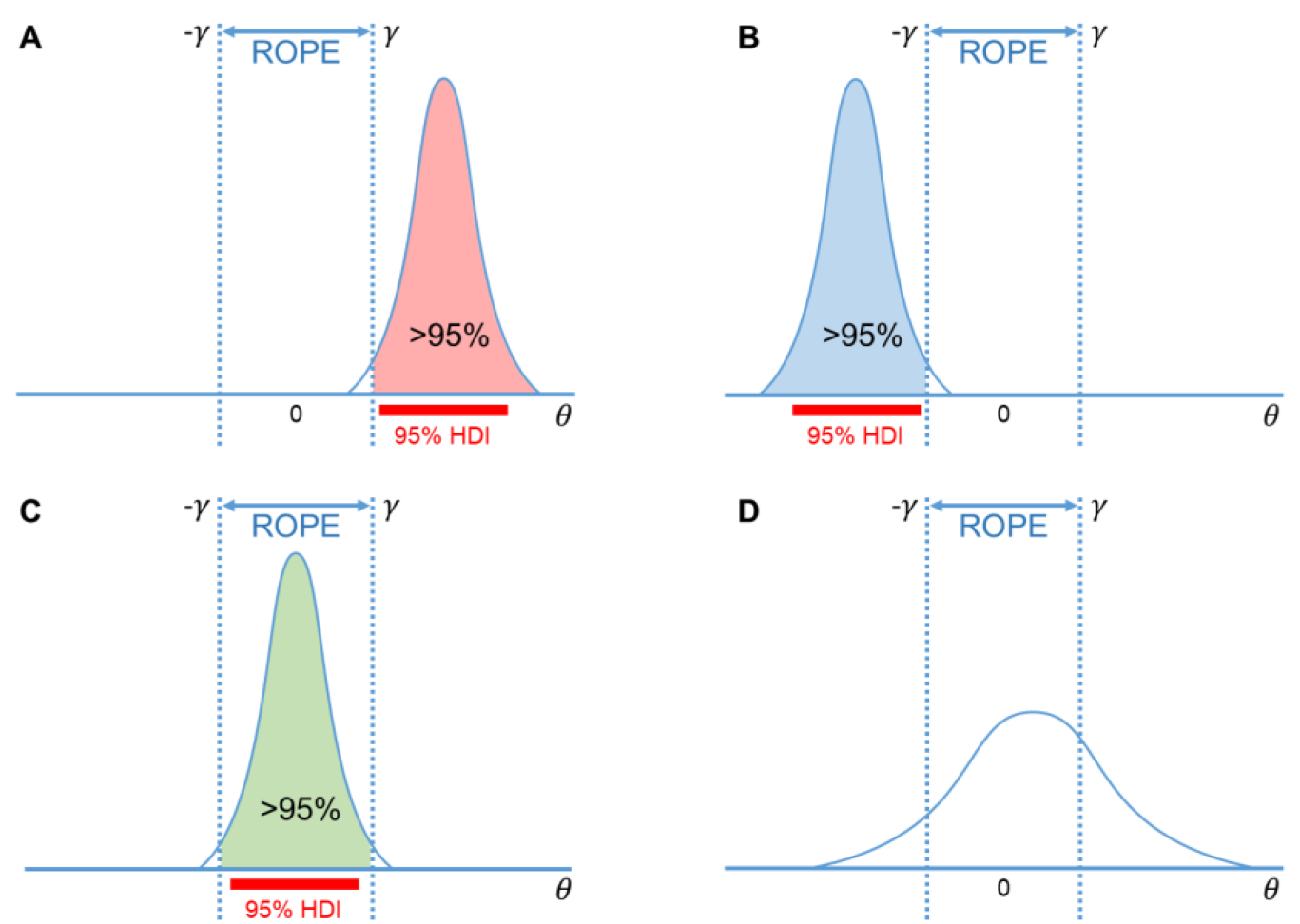
Possible variants of the posterior probability distributions of the effect *θ* = *cβ* in A) ‘activated’ voxels, B) ‘deactivated’ voxels, C) ‘not activated’ voxels, D) ‘low confidence’ voxels. The ‘ROPE only’ rule considers only the coloured parts of the distributions. The ‘HDI+ROPE’ rule considers overlap between the ROPE and 95% HDI. Scheme modified from Magerkurth et al. (2015)

### 2.6. Bayesian model inference

With BPI, we consider the posterior probabilities of the linear contrast of parameters *θ* = *cβ*. Instead, we can consider models using BMI.

Let *H_alt_* and *H_null_* be two non-overlapping hypotheses represented by models *M_alt_* and *M_null_*. These models are defined by two parameter spaces: 1) *M_alt_*: *θ* > *γ* and *θ* < −*γ*, and 2) *M_null_* −*γ* ≤ *θ* ≤ *γ*.

Now, we can rewrite Bayes’ rule (eq. 1) for *M_alt_* and *M_null_*

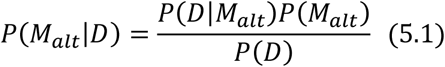

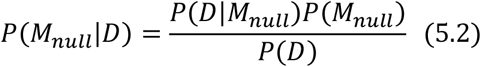

If we divide equation (5.1) by (5.2), *P*(*D*) is cancelled out, and we obtain:

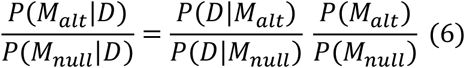

In verbal form equation (6) can be expressed as:

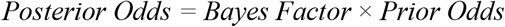

The Bayes factor (*BF*) is a multiplier that converts prior model probability odds to posterior model probability odds. It indicates the relative evidence for one model against another. For example, if 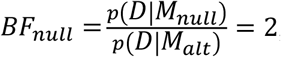, then the observed data are twice as likely under the null model than under the alternative.

A connection exists between the BPI (eq. 2–4), and BMI (eq. 6) (see Morey and Rouder, 2011; Liao et al., 2019):

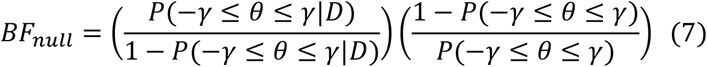

or, in verbal form:

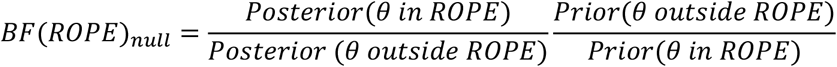

For convenience, *BF* may also be expressed in the form of a natural logarithm:

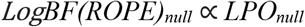

See Schematic illustration of *BF(ROPE)_null_* in Fig. 5A.

**Figure 5.**
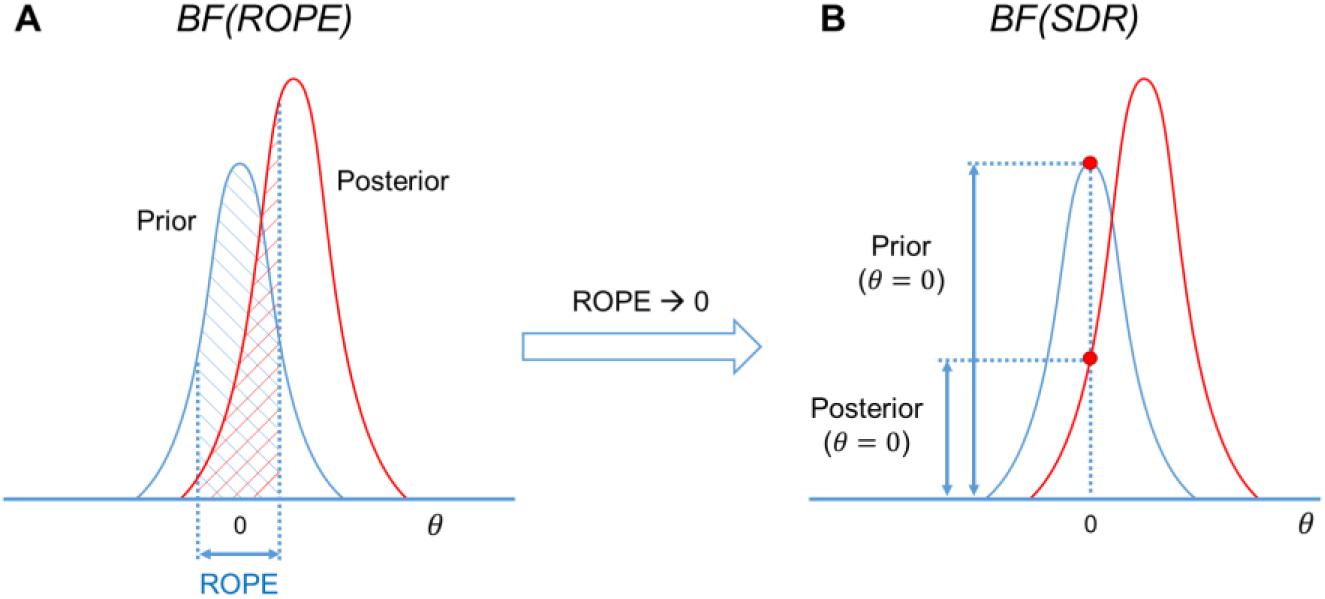
Schematic of *BFs* used in BMI. A) *BF(ROPE)* is related to the areas under the functions of the posterior and prior probability densities inside and outside the ROPE. B) *BF(SDR)* is the relation between the posterior and prior probability at *θ* = 0.

The calculation of *BF* may be computationally challenging, as it requires integration over the parameter spaces. However, if the ROPE has zero width (*γ* = 0), then the *BF* has an analytical solution known as the Savage–Dickey ratio (SDR) (Wagenmakers et al., 2010; Fritson and Penny, 2011; Rosa et al., 2012; Penny and Ridgway, 2013). The interpretation of the SDR is simple: if the effect size is less likely to equal zero after obtaining the data (posterior probability at *θ* = 0) than before (prior probability at *θ* = 0), then *BF(SDR)null* < 1; that is, we have more evidence for *Malt* (see Fig. 5B).

### 2.7. Relations between frequentist and Bayesian approaches

Now we can point out the conceptual links between the frequentist and Bayesian approaches.

1. **Parameter estimation.** When we have no prior information, that is, all parameter values are a priori equally probable (‘flat’ prior), Bayesian parameter estimation reduces to frequentist parameter estimation (maximum likelihood estimation; Friston et al., 2002a).
2. **Multiplicity adjustments.** One of the major concerns in frequentist inference is the multiplicity problem. In general, after the Bayesian parameter estimation, it is not necessary to classify any voxel as ‘(de)activated’ or ‘not activated’. If we consider *unthresholded* maps of posterior probabilities, *LPOs,* or *LogBFs,* the multiple comparisons problem does not arise (Friston and Penny, 2003). However, if we apply a decision rule to classify voxels, we should control for wrong decisions across multiple comparisons (Woolrich et al., 2009, see also possible loss functions in Muller et al., 2006; Kruschke and Liddell, 2017a). The advantage of hierarchical PEB with the ‘global shrinkage’ prior is that it automatically accounts for multiple comparisons without the need for ad hoc multiplicity adjustments (Berry, 1988; Friston and Penny, 2003; Scott and Berger, 2010; Gelman et al., 2012). The frequentist approach processes every voxel independently, whereas the PEB algorithm considers joint information from all voxels. Frequentist inference uncorrected for multiple independent comparisons is prone to label noise-driven, random extremes as ‘statistically significant’. Bayesian analysis specifies that extreme values are unlikely a priori, and thus they shrink toward a common mean (Lindley, 1990; Westfall et al., 1997; Berry and Hochberg, 1999; Friston et al., 2002a, 2002b; Gelman et al., 2012; Kruschke and Liddell, 2017b). If we consider *thresholded* maps of posterior probabilities, for example, *P_act_* > 95%, then as many as 5% of ‘activated’ voxels could be falsely labelled so. This is conceptually similar to the false discovery rate (FDR) correction (Berry and Hochberg, 1999; Frison et al., 2002b; Friston and Penny, 2003; Storey, 2003; Muller et al., 2006; Schwartzman et al., 2009). In practice, BPI with *γ* = 0 should produce similar results (in terms of the number of ‘(de)activated’ voxels) as classical NHST with FDR correction. If we increase the ES threshold, fewer voxels will be classified as ‘(de)activated’, and at some *γ* value, BPI will produce results similar to the more conservative Family Wise Error (FWE) correction^2^.
3. **Interval-based hypothesis testing**. Frequentist interval-based hypothesis testing is conceptually connected with BPI, particularly, the ‘HDI+ROPE’ decision rule. The former considers the intersection between ROPE and the confidence intervals. The latter considers the intersection between ROPE and the HDI (credible intervals).
4. **BPI and BMI.** BMI based on *BF(ROPE)* is conceptually linked to BPI based on the ‘ROPE-only’ decision rule. The interval-based Bayes factor *BF(ROPE)* is proportional to the posterior probability odds. When ROPE is infinitesimally narrow, *BF* can be approximated using the *SDR*. Note that even though *BF(SDR)* is based on the point-null hypothesis, it can still provide evidence for the null hypothesis, in contrast to BPI with *γ* = 0. However, *BF(SDR)* in PEB settings has not yet been tested using empirical fMRI data. Because the point-null hypothesis is always false (Meehl, 1967), BPI and *BF(ROPE)* are preferred over *BF(SDR)*.

### 2.8. Definition of the effect size threshold

The main difficulty in applying interval-based methods is the choice of the ES threshold *γ*. To date, only a few studies have been devoted to determining the minimal relevant effect size. One of them suggested a method to objectively define *γ* at the subject level of analysis which was calibrated by clinical experts and may be implemented for pre-surgical planning (Magerkurth et al., 2015). At the same time, the problem of choosing the ES threshold *γ* for group-level Bayesian analysis remains unresolved.

Several ways in which to define the ES threshold are available. Firstly, we can conduct a pilot study to determine the expected effect sizes. Secondly, we can use data from the literature to determine the typical effect sizes for the condition of interest. Thirdly, we can use the default ES thresholds that are commonly accepted in the field. One of the first ES thresholds proposed in the neuroimaging literature was *γ =* 0.1% (Friston et al., 2002b). This is the default ES threshold for the subject-level BPI in SPM12. For the group-level BPI, the default ES threshold is one prior standard deviation of the effect *γ* = 1 *prior SD_θ_* (Friston and Penny, 2003). Fourthly, γ can be selected in such a way as to ensure maximum similarity of the activation patterns revealed by classical NHST and Bayesian inference. This would allow us to reanalyse the data using Bayesian inference, reveal similar activation patterns as was previously the case for classic inference, and detect the ‘not activated’ and ‘low confidence’ voxels. Lastly, we can consider the posterior probabilities at multiple ES thresholds or compute the ROPE maps (see below).

It should also be noted that the ES threshold can be expressed as unstandardised (raw *β* values or percent signal change) and standardised values (for example, Cohen’s d). Raw *β* values calculated by SPM12 at the first level of analysis represent the BOLD signal in arbitrary units. However, they can be scaled to a more meaningful unit, the BOLD percent signal change (PSC) (Poldrack et al., 2011; Chen et al., 2017). Unstandardised and standardised values have disadvantages and advantages. Different ways exist in which to scale β values to PSC (Pernet, 2014; Chen et al., 2017), which is problematic when comparing the results of different studies. Standardised values represent the effect size in terms of the standard deviation units, which supposedly facilitate the comparison of results between different experiments. However, standardised values are relatively more unstable between measurements and less interpretable (Baguley, 2009; Chen et al., 2017). Moreover, Cohen’s d is closely related to the t-value (for one sample case, 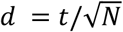) and may share some drawbacks with t-values. Reimold et al. (2005) showed that spatial smoothing has a nonlinear effect on voxel variance. Using t-values or Cohen’s d for inference in neuroimaging may lead to spatially inaccurate results (spatial shift of local maxima in t-maps or Cohen’s d maps compared to PSC-maps). In this study, we focused on PSCs.

## 3. Methods

We used the HCP and UCLA datasets to: 1) compare classical NHST and BPI, 2) consider different approaches to ES thresholding and estimate typical effect sizes, 3) demonstrate the behaviour of classical NHST and BPI depending on the sample size and spatial smoothing, and 4) illustrate a possible practical application of BPI.

### 3.1. Datasets

Seven block-design tasks were considered from the HCP dataset, including working memory, gambling, motor, language, social cognition, relation processing, and emotion processing tasks (Barch et al., 2013). Two event-related tasks, including the stop-signal and task-switching tasks were considered from the UCLA dataset (Poldrack et al., 2016). The length, conditions, and number of scans of the tasks are provided in the Supplementary Materials (Table S1). A subset of 100 unrelated subjects (S1200 release) was selected from the HCP dataset (54 females, 46 males, mean age = 29.1 ± 3.7 years) for assessment. A total of 115 subjects from the UCLA dataset were included in the analysis (55 females, 60 males, mean age = 31.7 ± 8.9 years) after removing subjects with no data for the stop-signal task, a high level (>15%) of errors in the Go-trials, and those of which the raw data were reported to be problematic (Gorgolewski et al., 2017). See the fMRI acquisition parameters in the Supplementary Materials, Par. 1.

### 3.2. Preprocessing

The minimal preprocessing pipelines for the HCP and UCLA datasets were described by Glasser et al. (2013) and Gorgolewski et al. (2017), respectively. Spatial smoothing was applied to the preprocessed images with a 4 mm full width at half maximum (FWHM) Gaussian smoothing kernel. Additionally, to compare the extent to which the performance of classical NHST and BPI depended on the smoothing, we applied 8 mm FWHM smoothing to the emotion processing task. Spatial smoothing was performed using SPM12. The results are reported for the 4 mm FWHM smoothing filter, unless otherwise specified.

### 3.3. Parameter estimation

Frequentist parameter estimation was applied at the subject level of analysis. A detailed description of the general linear models for each task design is available in the Supplementary Materials, Par. 2. Fixation blocks and null events were not modelled explicitly in any of the tasks. Twenty-four head motion regressors were included in each subject-level model (six head motion parameters, six head motion parameters one time point before, and 12 corresponding squared items) to minimise head motion artefacts (Friston et al., 1996). Raw *β* values were converted to PSC relative to the mean whole-brain ‘baseline’ signal (Supplementary Materials, Par. 3). The linear contrasts of the *β* values were calculated to describe the effects of interest *θ* = *cβ* in different tasks. The sum of positive terms in the contrast vector, *c*, is equal to one. The list of contrasts calculated in the current study to explore typical effect sizes is presented in Table S1. At the group level of analysis, the Bayesian parameter estimation with the ‘global shrinkage’ prior was applied using SPM12 (v6906). We performed a one-sample test on the linear contrasts created at the subject level of analysis.

### 3.4. Classical NHST and Bayesian parameter inference

Classical inference was performed using voxel-wise FWE correction with *α* = 0.05. This is the default SPM threshold and is known to be conservative and to guarantee protection from false positives (Eklund et al., 2016). Although voxel-wise FWE correction may be too conservative for small sample sizes, it is recommended when large sample sizes are available (Woo et al., 2014).

BPI, accessible via the SPM12 GUI, allows the user to declare only whether the voxels are ‘activated’ or ‘deactivated’. The classification of voxels as being either ‘not activated’ or ‘low confidence’ requires the posterior mean and variance. At the group level of analysis, SPM12 does not save the posterior variance image. However, the posterior variance can be reconstructed from the image of the noise hyperparameter using a first-order Taylor series approximation (Penny and Ridgway, 2013). Therefore, in the current study, BPI was performed using in-house scripts available at (https://github.com/Masharipov/Bayesian_inference). For the ‘ROPE-only’ rule, the posterior probability threshold was *P_thr_* = 95% (*LPO* > 3). For the ‘HDI+ROPE’ rule, we used 95% HDI.

We compared the number of ‘activated’ voxels (as a percentage of the total number of voxels) detected by Bayesian and classical inference. We also compared the number of ‘activated,’ ‘deactivated,’ and ‘not activated’ voxels detected using BPI with the ‘ROPE-only’ and ‘HDI+ROPE’ decision rules and different ES thresholds. To estimate the influence of the sample size on the results, all the above-mentioned analyses were performed with samples of different sizes: 5 to 100 subjects from the HCP dataset and 5 to 115 subjects from the UCLA dataset, in steps of 5 subjects. Ten random groups were sampled for each step.

### 3.5. Effect size thresholds

We considered three ES thresholds: firstly, the default ES threshold for the subject-level *γ* = 0.1% (BOLD PSC); secondly, the default ES threshold for the group-level *γ* = 1 *prior SD_θ_*; thirdly, the *γ*(*Dice_max_*) threshold, which ensures maximum similarity of the activation patterns revealed by classical NHST and BPI. The similarity was assessed using the Dice coefficient:

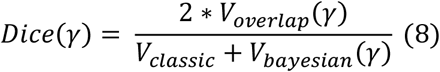

where *V_classic_* is the number of ‘activated’ voxels detected using classical NHST, *V_bayesian_*(*γ*) is the number of ‘activated’ voxels detected using BMI with the ES threshold *γ*, and *V_overlap_* is the number of ‘activated’ voxels detected by both methods. A Dice coefficient of 0 indicates no overlap between the patterns, and 1 indicates complete overlap. Dice coefficients were calculated for *γ* ranging from 0% to 0.4% in steps of 0.001%.

### 3.6. Estimation of typical effect sizes

In the current study, we aimed to provide a reference set of typical effect sizes for different task designs (block and event-related) and different contrasts (‘task-condition > control-condition,’ ‘task-condition > baseline,’) in a set of a priori defined regions of interest (ROI). Effect sizes were expressed in PSC and Cohen’s d. ROI masks were defined using anatomical and a priori functional masks. For more details, see Supplementary Materials, Par. 4.

### 3.7. Evaluating BPI on contrasts with no expected practically significant difference

BPI should be able to detect the ‘null effect’ in the majority of voxels when comparing samples with no expected practically significant difference. For example, there may be two groups of healthy adult subjects performing the same task or two sessions with the same task instructions. To test this, we used fMRI data from the emotion processing task. To emulate two ‘similar’ *independent* samples, 100 healthy adult subjects’ contrasts (‘Emotion > Shape’) were randomly divided into two groups of 50 subjects. A two-sample test comparing the ‘Group #1’ and ‘Group #2’ was performed with the assumption of unequal variances between the groups (SPM12 default option). To emulate ‘similar’ *dependent* samples, we randomized ‘Emotion > Shape’ contrasts from right-to-left (RL) and left-to-right (LR) phase encoding sessions in the ‘Session #1’ and ‘Session #2’ samples. Each sample consisted of 50 contrasts from the RL session and 50 from the LR session. A paired test designed to compare ‘Session #1’ and ‘Session #2’ was equivalent to the one-sample test on 50 ‘RL > LR session’ and 50 ‘LR > RL session’ contrasts.

## 4. Results

### 4.1. Results for contrasts with no expected practically significant difference

Classical NHST did not show a significant difference between ‘Group #1’ and ‘Group #2’ (see Supplementary Materials, Fig. S1). BPI with the ‘ROPE-only’ decision rule and default ES threshold *γ* = 1 *prior SD_θ_* = 0.190% classified 83.4% of voxels as having ‘no difference’ in which the null hypothesis was accepted (see Fig. S1). The ‘HDI+ROPE’ rule classified 76.2% of voxels as having ‘no difference’.

Classical NHST did not reveal a significant difference between ‘Session #1’ and ‘Session #2’ (see Fig. S2). The *prior SD_θ_* was 0.005%. In this case, using the default ES threshold *γ* = 1 *prior SD_θ_* did not allow the detection of any ‘no difference’ voxels, because the ROPE was unreasonably narrow. The ‘null effect’ was detected in all voxels beginning with a *γ =* 0.013% threshold using the ‘ROPE-only’ and ‘HDI+ROPE’ decision rules (see Fig. S2).

In this way, when comparing two ‘similar’ *independent* samples (two groups of healthy subjects performing the same task), BPI with the default group-level threshold (*one prior SD_θ_*) allowed us to correctly label voxels as having ‘no difference’ for the majority of the voxels of the brain. However, when comparing two ‘similar’ *dependent* samples (two sessions from the same task), the *one prior SD_θ_* threshold became inadequately small.

Therefore, the default *one prior SD_θ_* threshold is not suitable when the difference between *dependent* conditions is very small (paired sample test or one-sample test). In such cases, one can use an a priori defined ES threshold based on previously reported effect sizes or provide an ES threshold at which most of the voxels can be labelled as having ‘no difference’, allowing the critical reader to decide whether this speaks in favour of the absence of differences.

### 4.2. Comparison of classical NHST and BPI results

Generally, classical NHST with voxel-wise FWE correction and BPI with the ‘ROPE-only’ decision rule and default group-level ES threshold *γ* = 1 *prior SD_θ_* revealed similar (de)activation patterns in all considered contrasts (see Fig. 6, Table 1, Supplementary materials, Tables S2–S10). The median ES threshold based on *Dicemax* and median default group-level ES threshold across all considered contrasts were close in magnitude to the default subject-level ES threshold *γ* = 0.1%: *γ*(*Dice_max_*) = 0.118% and *γ* = 1 *prior SD_θ_* = 0.142%. The median *Dice_max_* across all the considered contrasts reached 0.904. At the same time, BPI allowed us to classify ‘non-significant’ voxels as ‘not activated’ or ‘low confidence’. As it can be clearly seen from Figure 6, areas with ‘non-activated’ voxels surround clusters with ‘(de)activated’ voxels. Both are separated by areas comprising ‘low confidence’ voxels.

**Figure 6.**
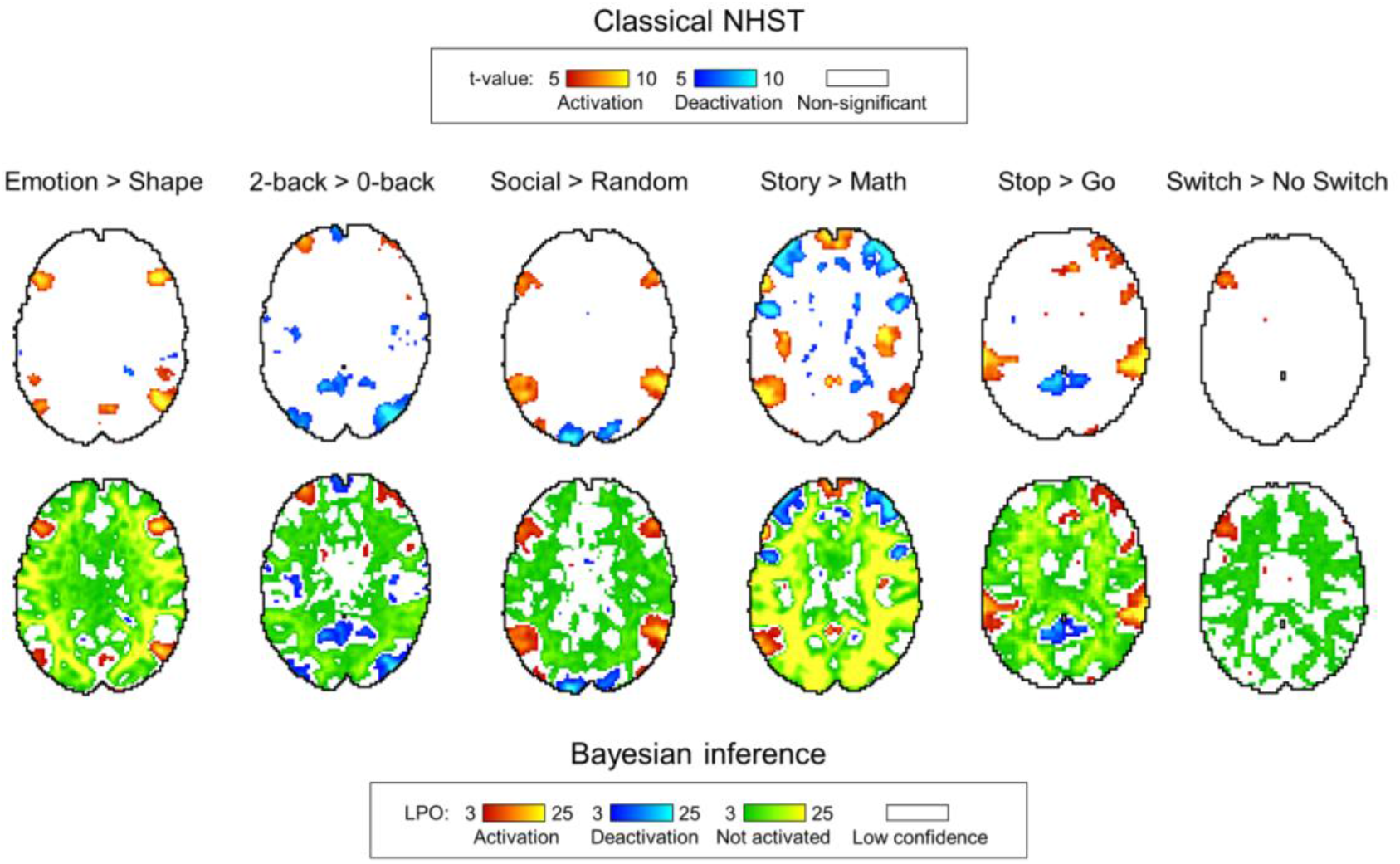
Examples of results obtained with classical NHST and BPI. Six contrasts were chosen for the illustration purposes (four event-related and two block-design tasks). Classical NHST was implemented using voxel-wise FWE correction (*α* = 0.05). BPI was implemented using the ‘ROPE-only’ decision rule, *P_thr_* = 95% (*LPO* > 3) and *γ* = 1 *prior SD_θ_*. Axial slice z = 18 mm (MNI152 standard space).

**Table 1.** Maximum Dice coefficient and corresponding effect size thresholds for each task.

| Contrast, $\theta$ | 1 prior $SD_{\theta}$, % | ‘ROPE-only’ decision rule | | ‘HDI+ROPE’ decision rule | |
| --- | --- | --- | --- | --- | --- |
| | | $\gamma(Dice_{max})$, % | $Dice_{max}$ | $\gamma(Dice_{max})$, % | Dice |
| Emotion processing |  |  |  |  |  |
| Emotion > Shape | 0.135 | 0.116 | 0.904 | 0.104 | 0.912 |
| Working memory |  |  |  |  |  |
| 2-back > baseline | 0.325 | 0.136 | 0.925 | 0.125 | 0.932 |
| 2-back > 0-back | 0.089 | 0.095 | 0.891 | 0.089 | 0.903 |
| Language |  |  |  |  |  |
| Story > Math | 0.255 | 0.119 | 0.896 | 0.108 | 0.904 |
| **Motor** |  |  |  |  |  |
| LF > baseline | 0.149 | 0.148 | 0.897 | 0.135 | 0.907 |
| RF > baseline | 0.171 | 0.160 | 0.886 | 0.144 | 0.897 |
| T > baseline | 0.268 | 0.205 | 0.904 | 0.181 | 0.913 |
| **Gambling** |  |  |  |  |  |
| Reward > baseline | 0.254 | 0.132 | 0.917 | 0.122 | 0.924 |
| Loss > baseline | 0.249 | 0.134 | 0.918 | 0.118 | 0.925 |
| Reward > Loss | 0.032 | 0.044 | 0.894 | 0.037 | 0.886 |
| **Social cognition** |  |  |  |  |  |
| Social > baseline | 0.325 | 0.139 | 0.939 | 0.124 | 0.944 |
| Social > Random | 0.104 | 0.114 | 0.896 | 0.104 | 0.907 |
| **Relational processing** |  |  |  |  |  |
| Relational > baseline | 0.390 | 0.154 | 0.935 | 0.143 | 0.940 |
| Relational > Match | 0.051 | 0.073 | 0.892 | 0.066 | 0.894 |
| **Stop-signal task** |  |  |  |  |  |
| Correct Stop > baseline | 0.069 | 0.066 | 0.895 | 0.061 | 0.906 |
| Correct Stop > Go | 0.064 | 0.052 | 0.906 | 0.047 | 0.917 |
| **Task-switching** |  |  |  |  |  |
| Switch > baseline | 0.133 | 0.075 | 0.907 | 0.067 | 0.916 |
| Switch > No switch | 0.030 | 0.037 | 0.924 | 0.033 | 0.925 |
| **Summary** |  |  |  |  |  |
| Median | 0.142 | 0.118 | 0.904 | 0.106 | 0.913 |

As expected, compared with the ‘HDI+ROPE’ rule, using the ‘ROPE-only’ rule slightly increases the number of ‘(de)activated’ and ‘not activated’ voxels (see Table 1 and Tables S2-10). The ‘HDI+ROPE’ rule labelled more voxels as ‘low confidence’. In principle, the difference between these rules would only be expected to increase with skewed distributions, which is not the case in SPM12. Thus, we recommend using the ‘ROPE-only’ decision rule (as well as in Friston et al., 2002a, 2002b, 2003; Wellek, 2010; Liao et al., 2019). In the following sections, we focus primarily on describing the results for the ‘ROPE-only’ rule.

### 4.3. Comparison of BPI results with different ES thresholds

Here, we first consider the results for the emotional processing task and then consider other tasks. Using the default single-subject ES threshold *γ* = 0.1% for the emotional processing task (‘Emotion > Shape’ contrast), 58.8% of all voxels can be classified as ‘not activated,’ 30.8% as ‘low confidence,’ and 10.1% as ‘activated’ (see Fig. 7, Table S2). The default group-level ES threshold *γ* = 1 *prior SD_θ_* = 0.135% allowed us to classify 75.0% of all voxels as ‘non-activated,’ 17.5% as ‘low confidence,’ and 7.4% as ‘activated’ (see Fig. 7, Table S2). Both types of thresholds were comparable to those of classical NHST for the detection of ‘activated’ voxels. The maximum overlap between ‘activations’ patterns revealed by classical NHST and BPI was observed at *γ*(*Dice_max_*) = 0.116% (see Fig. 8, Table 1).

**Figure 7.**
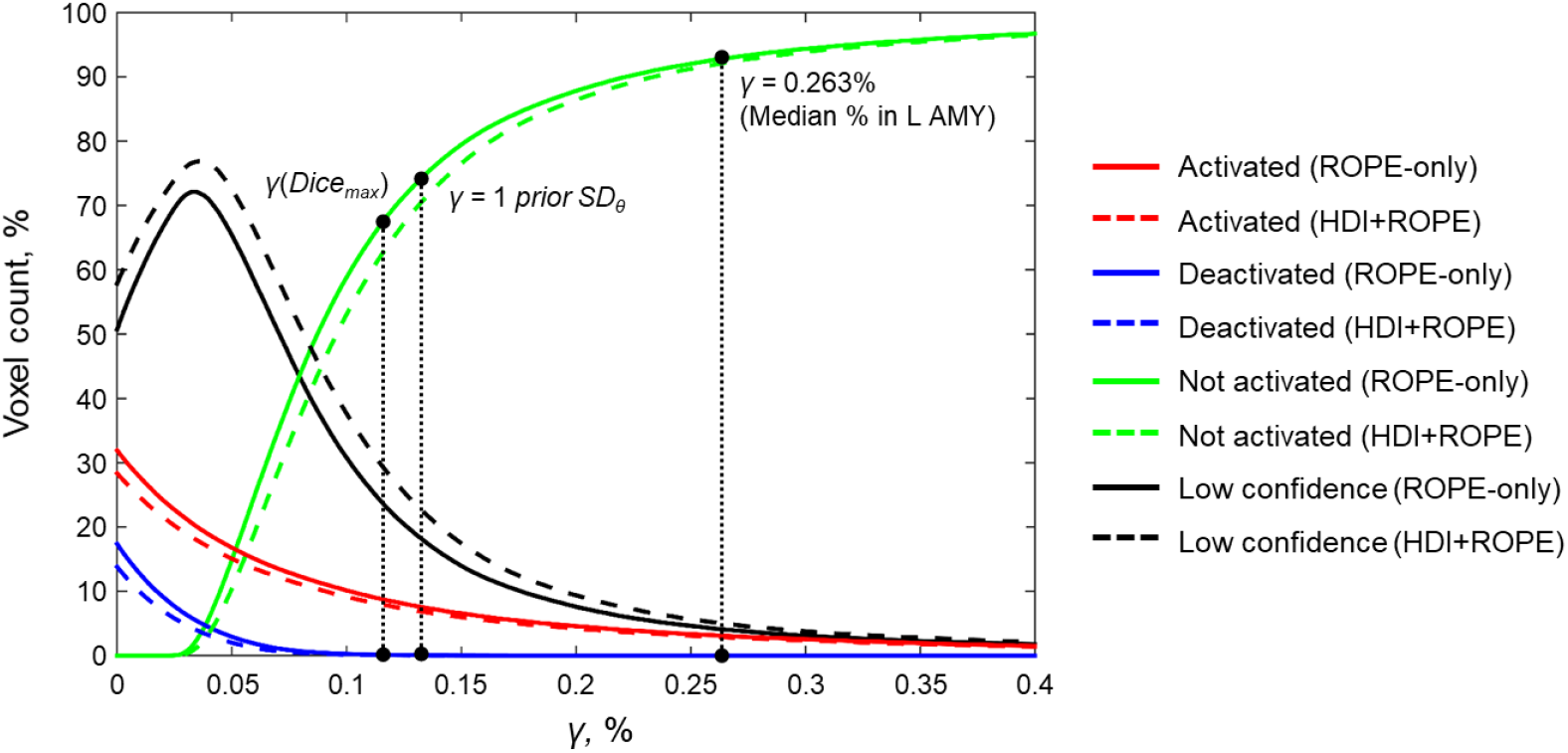
Number of voxels classified into the four categories depending on the ES threshold *γ*. The results for the emotion processing task (‘Emotion>Shape’ contrast) are presented for illustration. Abbreviations: L AMY – left amygdala.

**Figure 8.**
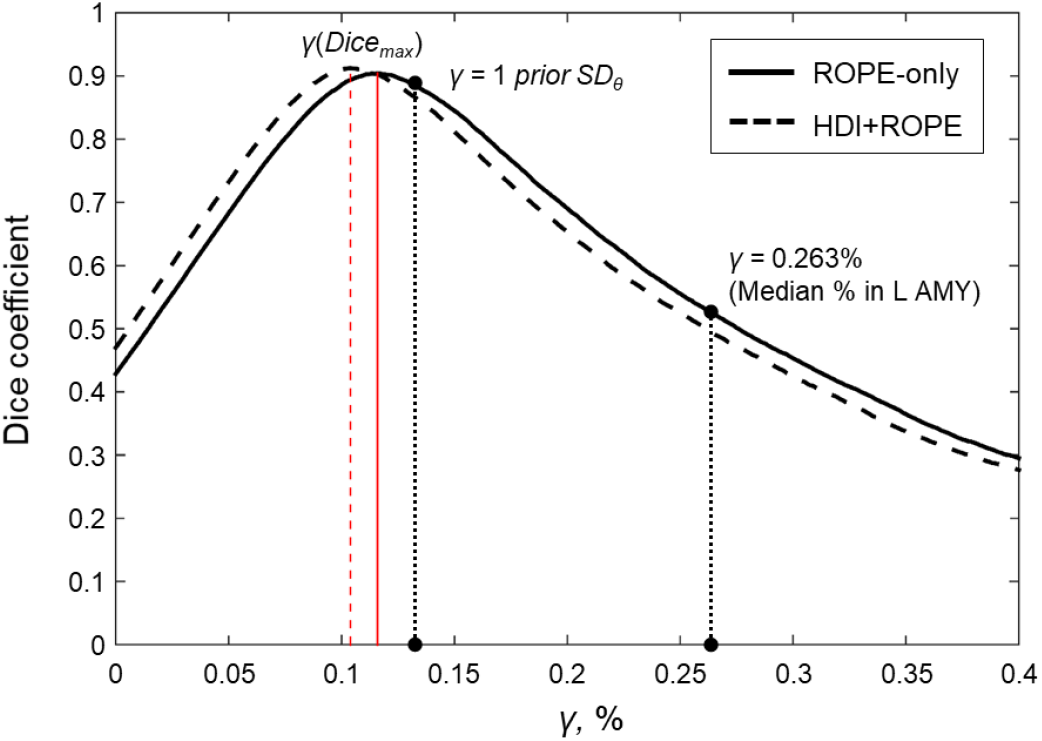
Dependence of the Dice coefficient on the ES threshold *γ*. Results for the emotion processing task (‘Emotion>Shape’ contrast). The red lines denote *γ*(*Dice_max_*). Abbreviations: L AMY – left amygdala.

For the ‘2-back > 0-back,’ ‘Left Finger > baseline,’ ‘Right Finger > baseline,’ and ‘Social > Random’ contrasts, the three ES thresholds that were considered—0.1%, *one prior SD_θ_, γ*(*Dice_max_*)—produced similar results (see Table 1 and Tables S3, S5, S7). For the event-related stop-signal task (‘Correct Stop > baseline’ and ‘Correct Stop > Go’ contrasts), *one prior SD_θ_* and *γ*(*Dice_max_*) were close in terms of their values but smaller than 0.1% (see Table 1). Block designs tend to evoke higher BOLD PSC than event-related designs; therefore, a lower *prior SD_θ_* should be expected for event-related designs and higher *prior SD_θ_* for block designs. Within a single design, in contrasts such as ‘task-condition > baseline’, higher BOLD PSC and *prior SD_θ_* would be expected than in contrasts in which the experimental conditions are compared directly. For example, the contrast ‘2-back > baseline’ has *prior SD_θ_* = 0.325% and contrast ‘2-back > 0-back’ has *prior SD_θ_* = 0.089%.

As previously noted, some contrasts did not elicit robust activations: ‘Reward > Loss’, ‘Relational > Match’, (Barch et al., 2013) and ‘Switch > No switch’ (Gorgolewski et al., 2017). The corresponding *γ*(*Dice_max_*) thresholds were 0.044%, 0.073%, and 0.037% (see Tables 1, S6, S8, and S10). The *prior SD_θ_* were 0.032%, 0.051%, and 0.030%. Correspondingly, BPI with the *γ* = 1 *prior SD_θ_* threshold classified 0%, 18.4%, and 42.2% of voxels as ‘not activated’. This demonstrates that when we compare conditions with similar neural activity and minor differences, it becomes more difficult to separate ‘(de)activations’ from the ‘null effects’ using the *γ* = 1 *prior SD_θ_* threshold.

### 4.4. Typical effect sizes in fMRI studies

A complete list of effect sizes (BOLD PSC and Cohen’s d) estimated for different tasks and a priori defined ROIs is presented in the Supplementary Materials (Tables S11–19). Here, we focus only on the BOLD PSC. The violin plots for some of these are shown in Figure 9.

**Figure 9.**
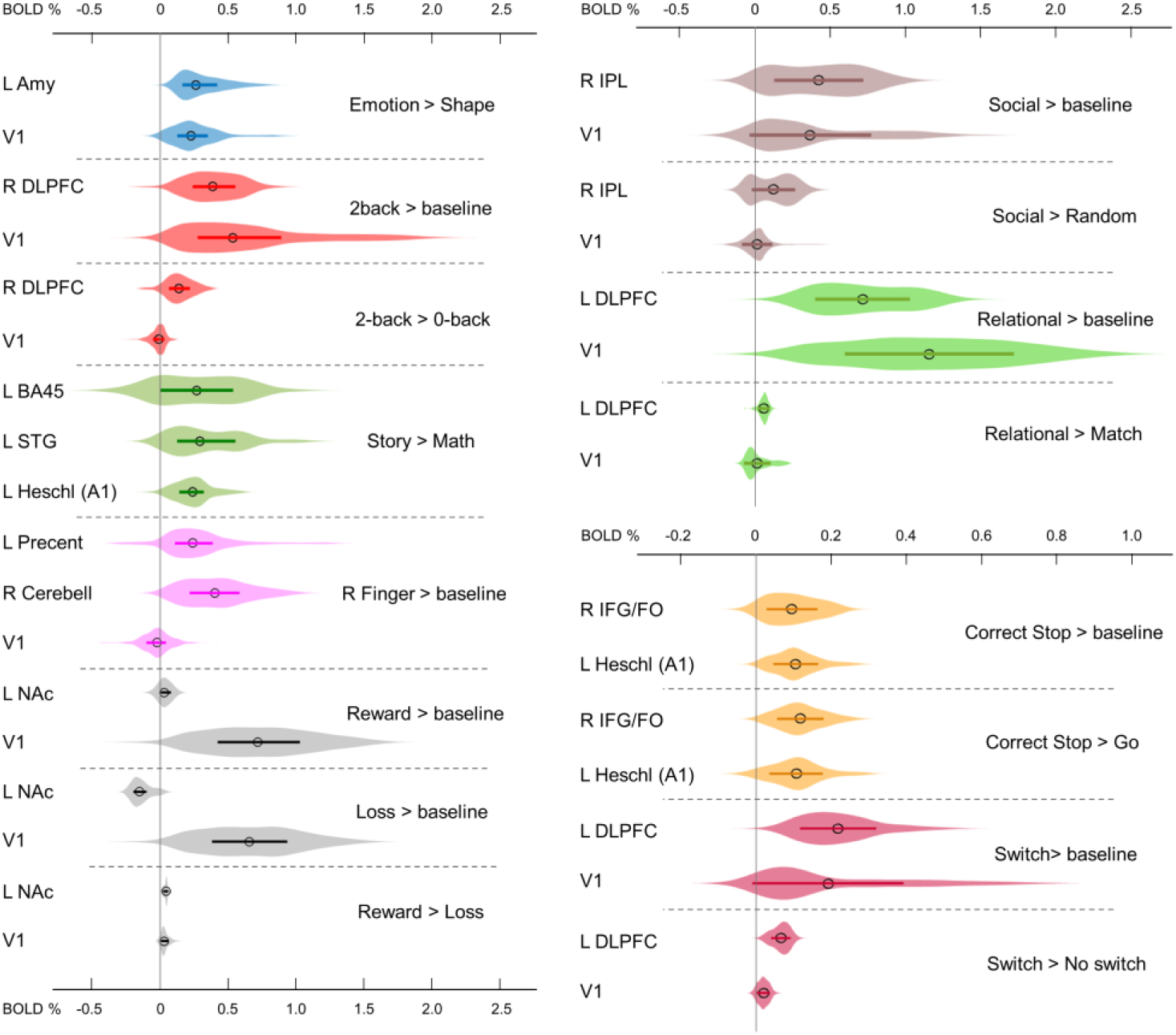
Typical BOLD PSC in fMRI studies. The box plots inside the violins represent the first and third quartile, and the black circles represent median values. Contrasts from the same task are indicated in one colour. Abbreviations: L/R – left/right, AMY – amygdala, V1 – primary visual cortex, DLPFC – dorsolateral prefrontal cortex, BA – Brodmann area, STG – superior temporal gyrus, A1 – primary auditory cortex, NAc – nucleus accumbens, IPL – inferior parietal lobule, IFG/FO – inferior frontal gyrus/frontal operculum.

For example, the median BOLD PSC in the left amygdala ROI, one of the key brain areas for emotional processing, was 0.263%, which is approximately twice as large as *one prior SD_θ_* (see Fig. 7). Thus, using this PSC as the ES threshold in future studies may cause the ROPE to become too wide compared to the effect sizes typical for tasks with such designs. Therefore, such a threshold can be used to detect large and highly localised effects. However, it may fail to detect small but widely distributed effects previously described for HCP data (Cremers et al., 2018).

In general, median PSCs within ROIs were up to 1% for block designs and 0.5% for event-related designs. The maximum PSCs reached 2.5% and were usually observed in the primary visual cortex (V1) for visual tasks comparing experimental conditions with baseline activity. For ‘moderate’ physiological effects, PSC varied in the range 0.1–0.2%, for example, for the ‘2-back > 0-back’ contrast, the median PSC in the right dorsolateral prefrontal cortex (R DLPFC in Fig. 9) was 0.137%. Likewise, for the ‘Social > Random’ contrast, the right inferior parietal lobule (R IPL) median PSC was 0.137%, for the ‘Correct Stop > Go’, the right inferior frontal gyrus/frontal operculum (R IFG/FO) median PSC was 0.120%. For more ‘strong’ physiological effects, the PSC was in the range 0.2–0.3%, for example, for the ‘Emotion > Shape’ contrast, the median PSC in the left amygdala was 0.263%, and for the ‘Story > Math’ contrast, the median PSC in the left Brodmann area 45 (Broca’s area) was 0.269%. For the motor activity, for example the ‘Right Finger > baseline’ contrast, the median PSC in the left precentral gyrus was 0.239%, in the left postcentral gyrus was 0.362%, in the left putamen was 0.290%, and in the right cerebellum was 0.401%. For the contrasts that did not elicit robust activations (Barch et al., 2013), the PSC was approximately 0.05%; for example, for the ‘Reward > Loss’ contrast, the median PSC in the left nucleus accumbens was 0.043%, and for the ‘Relational > Match’ contrast, the median PSC in the left dorsolateral prefrontal cortex was 0.062%.

### 4.5. ROPE maps

We considered BPI with two consecutive thresholding steps: 1) calculate the *LPOs* (or PPMs) with a selected ES threshold *γ*, 2) apply the posterior probability threshold *p_th_* = 95% or consider the overlap between 95% HDI and ROPE. We can *reverse the thresholding sequence* and calculate *the ROPE maps*.

For the ‘(de)activated’ voxels, the ROPE map contains the maximum ES thresholds that allow voxels to be classified as ‘(de)activated’ based on the ‘ROPE-only’ or ‘HDI+ROPE’ decision rules. For the ‘not activated’ voxels, the map contains the minimum effect size thresholds that allow voxels to be classified as ‘not activated’.

The procedure for calculating the ROPE map can be performed as follows. Let us consider a gradual increase in the ROPE radius (i.e., the half-width of ROPE or the ES threshold *γ*) from zero to the maximum effect size in observed volume. (1) For voxels in which PSC is close to zero, at a certain ROPE radius, the posterior probability of finding the effect within the ROPE becomes higher than 95%. This width is indicated on the ROPE map for ‘not activated’ voxels. (2) For voxels in which the PSC deviates from zero, at a certain ROPE radius, the posterior probability of finding the effect outside the ROPE becomes lower than 95%. This width is indicated on the ROPE map for ‘(de)activated’ voxels. The same maps can be calculated for the ‘HDI+ROPE’ decision rule.

Examples of the ROPE maps are shown in Figure 10. From our point of view, ROPE maps, as well as unstandardised effect size (PSC) maps, may facilitate an intuitive understanding of the spatial distribution of a physiological effect under investigation (Chen et al., 2017). They can also be a useful addition to standard PPMs.

**Figure 10.**
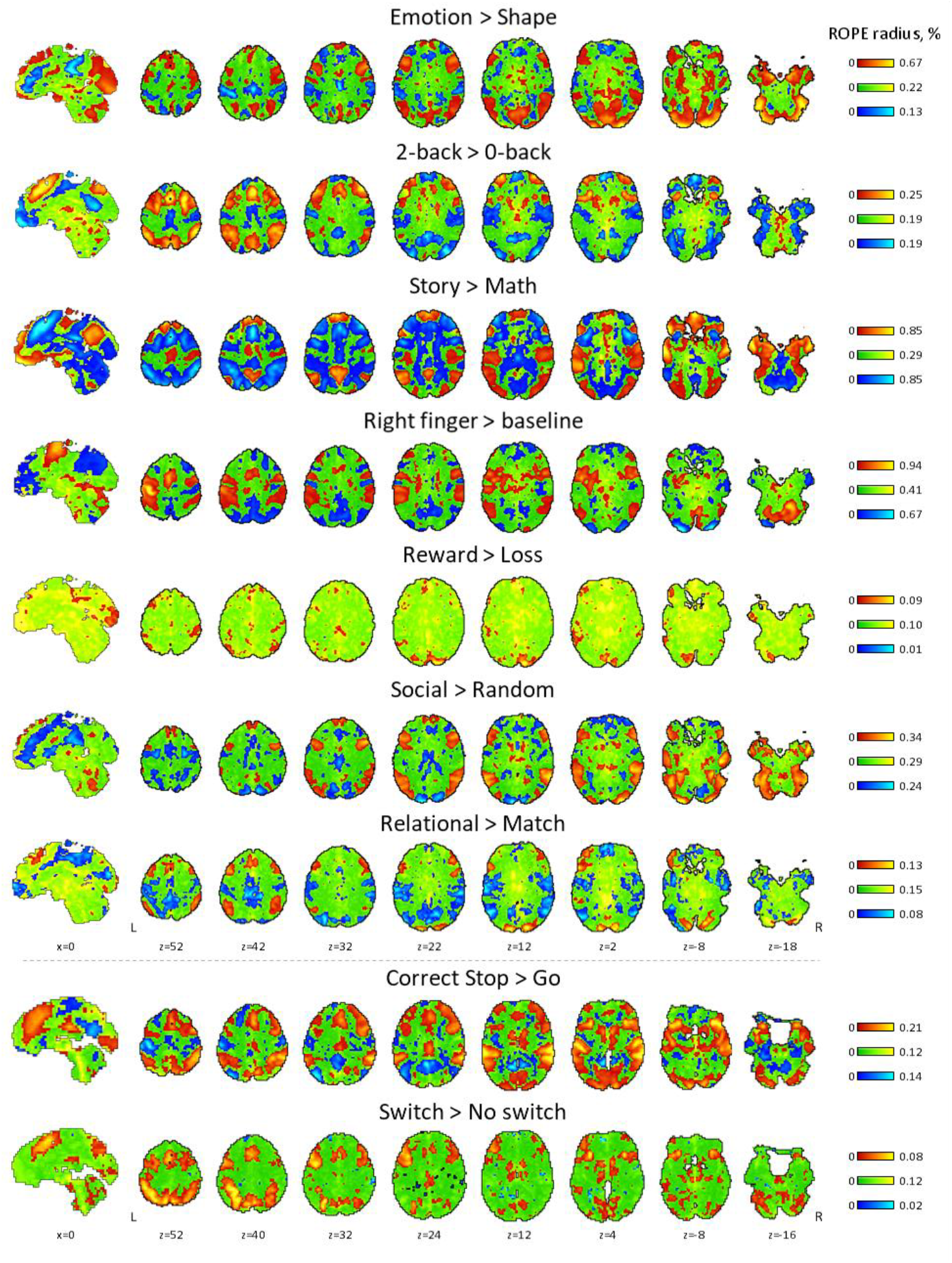
ROPE maps for all tasks considered in the present study. The ROPE maps are presented using different colours for the ‘activated’, ‘deactivated’, and ‘not activated’ voxels. The green bars represent the minimum ROPE radii at which voxels with a PSC close to zero can be classified as ‘not activated’ based on the ‘ROPE-only’ decision rule. The red and blue bars represent the maximum ROPE radii at which voxels of which the PSC deviates from zero can be classified as ‘activated’ and ‘deactivated’, respectively.

### 4.6. Effects of spatial smoothing on classical NHST and BPI

Two main differences were identified with respect to the influence of the spatial smoothing between classical NHST and BPI. Firstly, some voxels with a negligible BOLD PSC could be classified as ‘not significant’ at lower smoothing and as ‘activated’ at higher smoothing using classical NHST. At the same time, BPI classified these voxels as ‘low confidence’ at lower smoothing and as ‘not activated’ at higher smoothing. Higher spatial smoothing increased the number of both ‘(de)activated’ and ‘not activated’ voxels classified by BPI, and decrease the number of ‘low confidence’ voxels.

Secondly, higher smoothing blurred the spatial localisation of local maxima of t-maps and PPMs (*LPO*-maps) to a different extent. Consider, for example, the emotion processing task (‘Emotion > Shape’ contrast). The broadening of two peaks in the left and right amygdala was more noticeable on the t-map than on the PPM (see Fig. 11).

**Figure 11.**
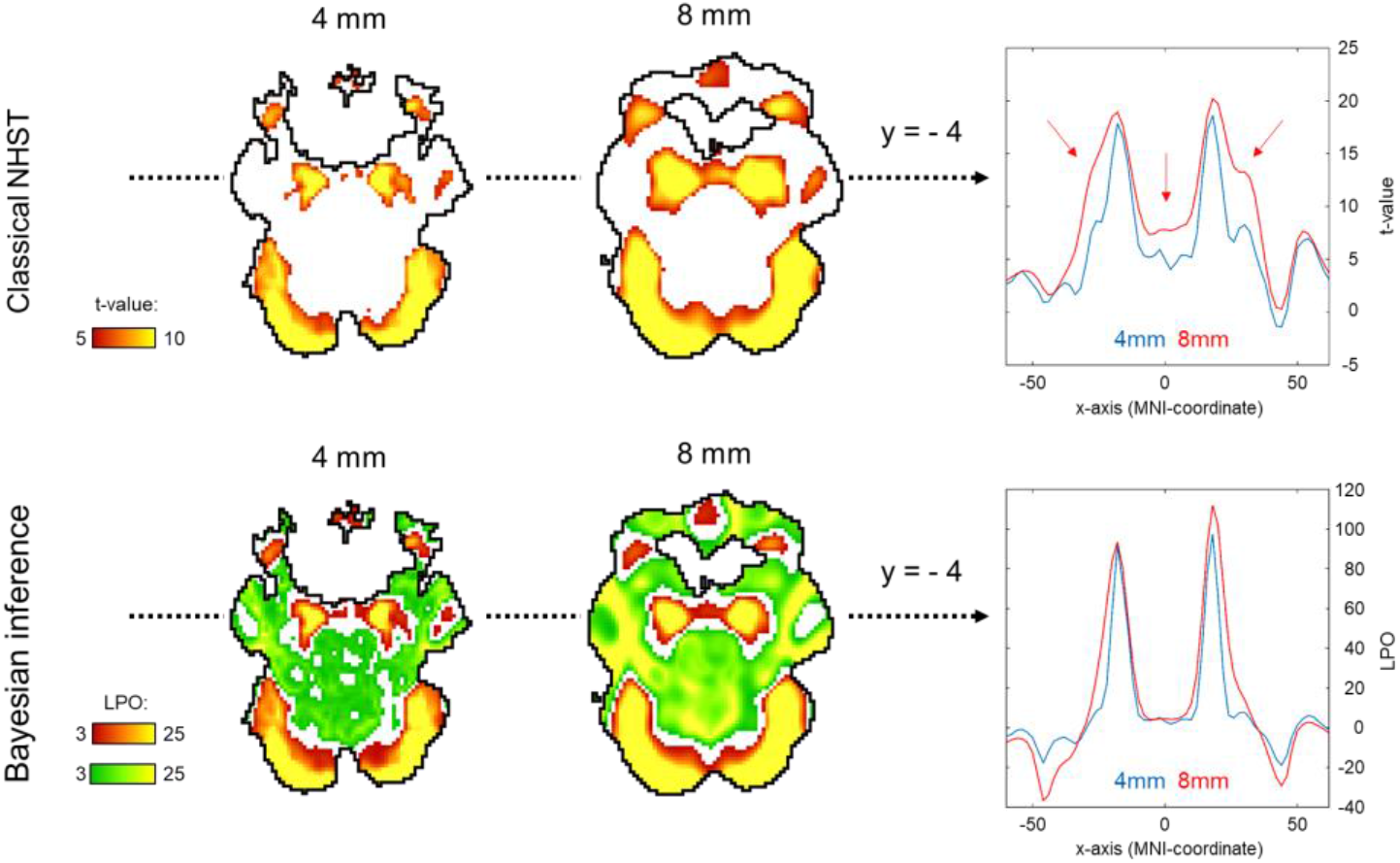
Influence of spatial smoothing on classical NHST and BPI: results for the emotion processing task (‘Emotion > Shape’ contrast). Classical NHST was implemented using voxel-wise FWE correction (*α* = 0.05). BPI was implemented using the ‘ROPE-only’ decision rule, *P_thr_* = 95% (*LPO* > 3) and *γ =* 1 *prior SD_θ_* = 0.135%. Axial slice z = –14 mm (MNI152 standard space). In the panels on the right, 1-D images are presented for t-values and *LPOs* along the x-axis for y = –4 mm. The red arrows indicate a noticeable broadening of two peaks of local maxima (left and right amygdala) at higher smoothing.

Smoothing was previously shown to have a nonlinear effect on the voxel variances and thus to affect more t-maps than *β* value maps, sometimes leading to counterintuitive artefacts (Reimold et al., 2005). This is especially noticeable at the border between two different tissues or between the two narrow peaks of the local maxima. If the peak is localised close to white matter voxels with low variability, then smoothing can shift the peak to the white matter. If low-variance white matter voxels separate two close peaks, then after smoothing, they may serve as a ‘bridge’ between the two peaks. To avoid this problem, Reimold et al. (2005) recommended using masked *β* value maps. In the present study, we suggest that PPMs based on BOLD PSC thresholding can mitigate this problem. Importantly, smoothing artefacts can also arise on Cohen’s d maps. Therefore, PPMs based on PSC thresholding may be preferable to PPMs based on Cohen’s d thresholding.

### 4.7. Sample size dependencies for classical NHST and BPI

An enlargement of the sample size led to an increase in the number of ‘activated’ and ‘not activated’ voxels, and a decrease in the number of ‘low confidence’ voxels. This is due to a decrease in the posterior variance. The curve of the ‘activated’ voxels rose much slower than that of the ‘not activated’ voxels. For the emotion processing task (‘Emotion > Shape’ contrast, block-design, two sessions, 352 scans), the largest gain in the number of ‘activated’ and ‘not activated’ voxels can be noted from 20 to 30 subjects (see Fig. 12A). With a sample size of *N* > 30, the number of ‘activated’ and ‘not activated’ voxels increased less steeply. The ‘not activated’ and ‘low confidence’ voxels curves intersected at *N* = 30 subjects. After the intersection point, the graphs reached a plateau.

**Figure 12.**
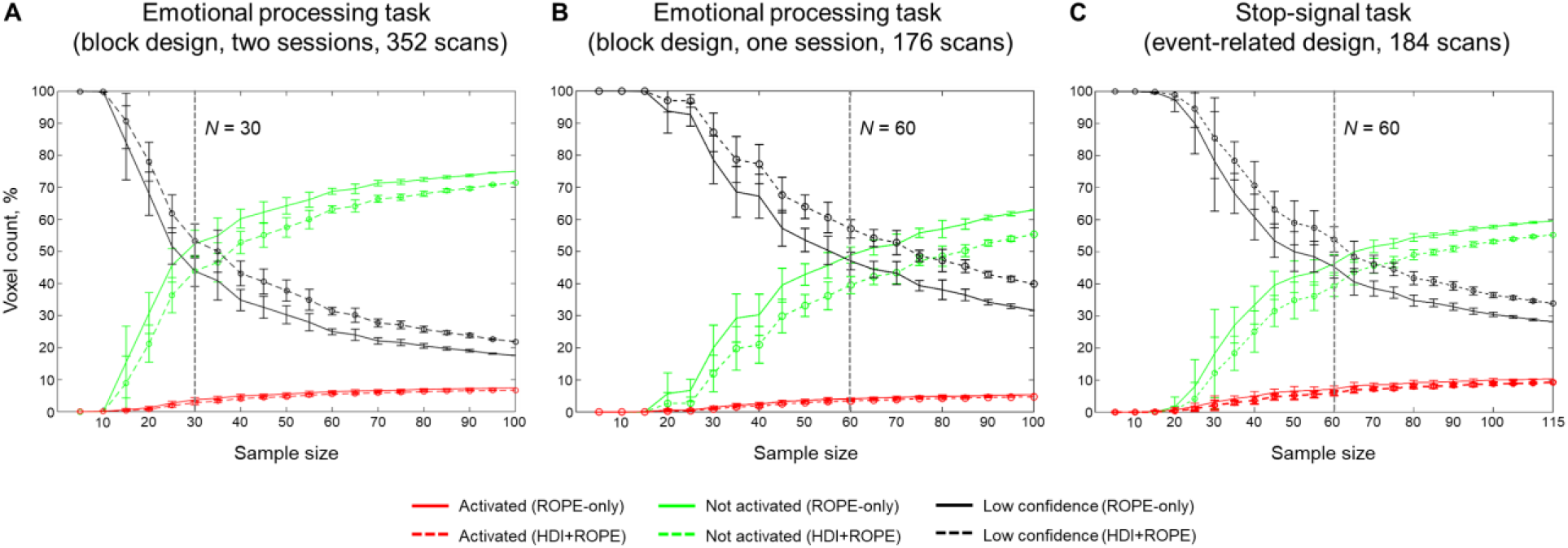
Dependencies of the number of ‘activated’, ‘not activated’, and ‘low confidence’ voxels on the sample size. BPI was implemented using *γ* = 1 *prior SD_θ_*. A) The emotional processing task (‘Emotion > Shape’ contrast, two sessions). B) The emotional processing task (‘Emotion > Shape’ contrast, one session). C) The stop-signal task (‘Correct Stop > Go’ contrast). The error bars represent the mean and standard deviation across ten random groups.

Considering only half of the emotional processing task data (one session, 176 scans), the intersection point shifted from *N* = 30 to *N* = 60 (see Fig. 12B). For the event-related task (‘Correct Stop > Go’ contrast, the stop-signal task, 184 scans), all considered dependencies had the same features as for the block-design task, and the point of intersection was at *N* = 60 subjects (see Fig. 12C). Therefore, the moment at which the graphs reach a plateau depends mainly on the amount of data at the subject level, that is, on the number of scans, blocks, and events. The task designs from the HCP and UCLA datasets have relatively short durations (for example, the stop-signal task has approximately 15 ‘Correct Stop’ trials per subject). Studies with a longer scanning time generally require a smaller sample size to enable inferences to be made with confidence.

Classical NHST with the voxel-wise FWE correction showed a steady linear increase in the number of ‘activated’ voxels with increasing sample size (see Fig. 13). With a further increase in the sample size, the number of statistically significant voxels revealed by classical NHST is expected to approach 100% (see, for example, Gonzalez-Castillo et al., 2012). In contrast, the BPI with the *γ* = 1 *prior SD_θ_* threshold demonstrated exponential dependencies. We observed a steeper increase at small and moderate sample sizes (*N* = 15–60). The curve of the ‘activated’ voxels flattened at large sample sizes (N > 80). BPI offers protection against the detection of ‘trivial’ effects that can appear as a result of an increased sample size if classical NHST with the point-null hypothesis is used (Friston et al., 2002a; Friston, 2012; Chen et al., 2017). This is achieved by the ES threshold *γ*, which eliminates physiologically (practically) negligible effects. Figure 13 presents an illustration of the Jeffreys-Lindley paradox, that is, the discrepancy between results obtained using classical and Bayesian inference, which is usually manifested at higher sample sizes (Jeffreys, 1939/1948; Lindley, 1957; Friston, 2012).

**Figure 13.**
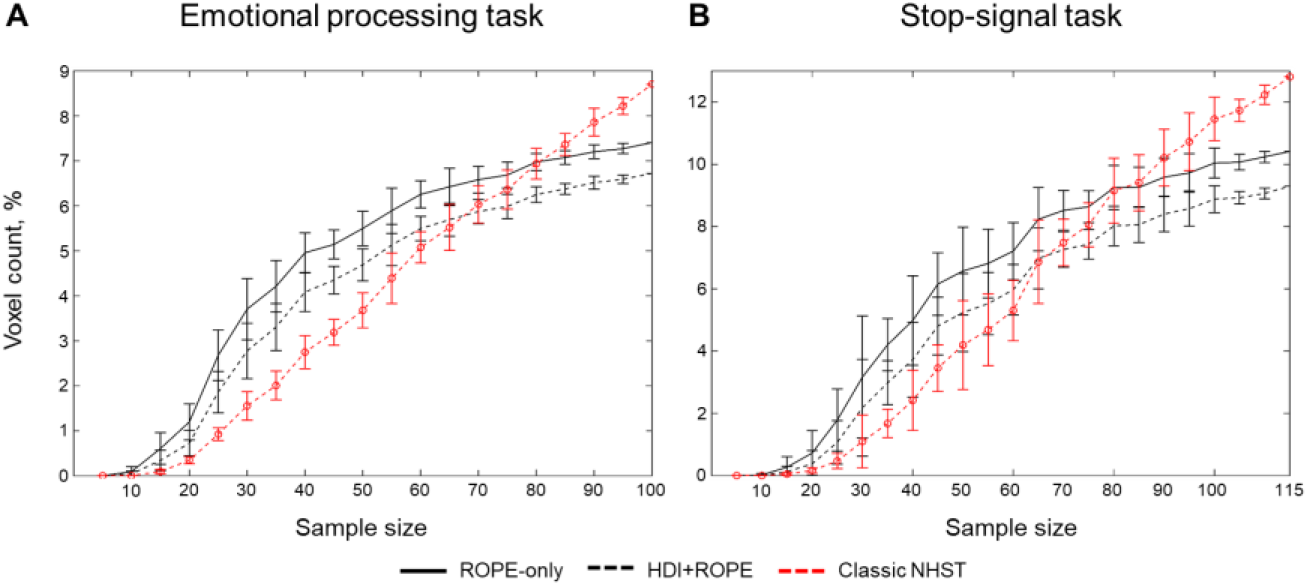
Dependencies of the number of ‘activated’ voxels on the sample size. Classical NHST was implemented using FWE correction (*α* = 0.05). BPI was implemented using *γ* = 1 *prior SD_θ_*. A) The emotional processing task (block design, ‘Emotion > Shape’ contrast). B) The stop-signal task (event-related design, ‘Correct Stop > Go’ contrast). The error bars represent the mean and standard deviation across ten random groups.

### 4.8. Example of practical application of BPI

In contrast to classical NHST, Bayesian inference allows us to:

1. Provide evidence that there is no practically meaningful BOLD signal change in the brain area when comparing the two task conditions.
2. Establish double dissociations (Friston et al., 2002a).
3. Provide evidence for practically equivalent engagement of one area under different experimental conditions in terms of local brain activity.
4. Provide evidence for the absence of a practically meaningful difference in BOLD signals between groups of subjects or repeated measures.

To illustrate a possible application of Bayesian inference in research practice, we used a working memory task. Let us consider an overlap between the ‘2-back > baseline’ and ‘0-back > baseline’ contrasts (see Fig 14, purple areas). We cannot claim that brain areas revealed by this conjunction analysis were equally engaged in the ‘2-back’ and ‘0-back’ conditions. To provide evidence for this notion, we can use BPI and attempt to identify voxels with a practically equivalent BOLD signal in the ‘2-back’ and ‘0-back’ conditions (see Fig 14, green). Overlap between the ‘2-back > baseline’ and ‘0-back > baseline’ and the ‘2-back = 0-back’ effects was found in several brain areas: visual cortex (V1, V2, V3), frontal eye field (FEF), superior eye field (SEF), parietal eye field (PEF, or posterior parietal cortex), lateral geniculate nucleus (LGN) and left primary motor cortex (M1) (see Fig. 14, white). This result can be easily explained by the fact that both experimental conditions require the subject to analyse perceptually similar visual stimuli and push response buttons with the right hand, which should not depend much on the working memory load. At the same time, it does not follow directly from simple conjunction analysis.

**Figure 14.**
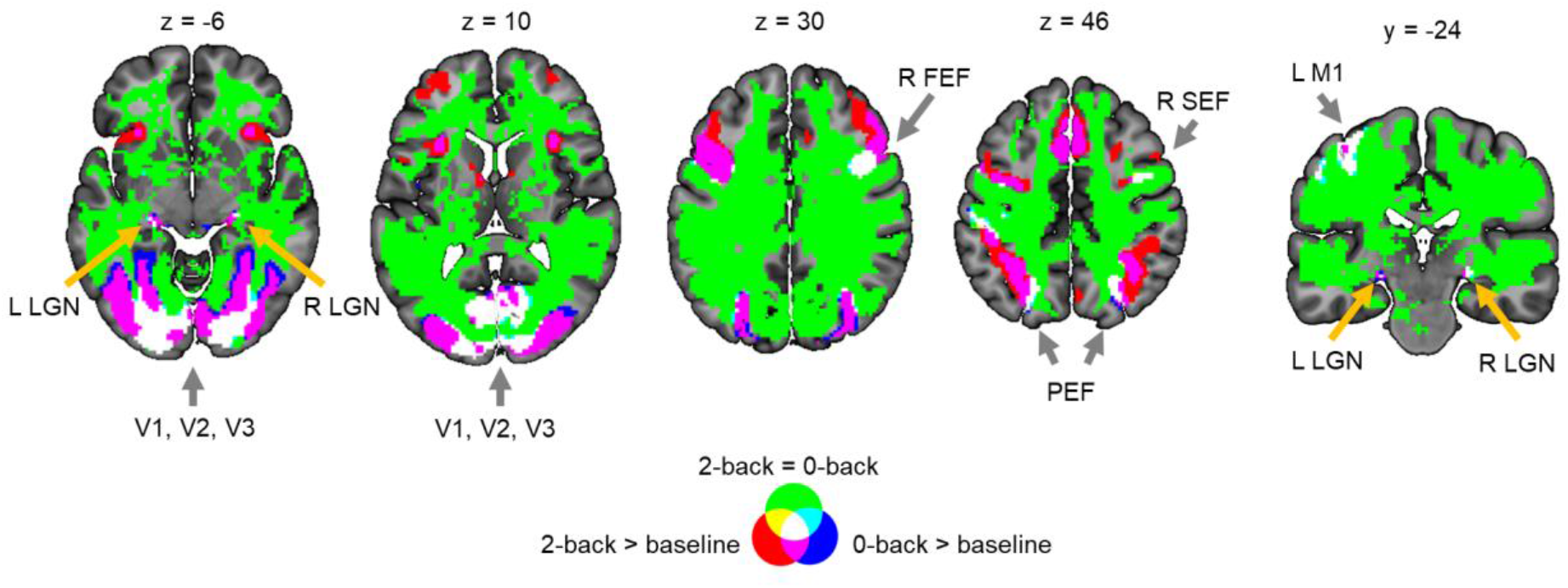
Example of possible application of BPI based on the working memory task. Abbreviations: L/R – left/right, V1, V2, V3 – primary, secondary, and third visual cortex, FEF – frontal eye field, SEF – superior eye field, PEF – parietal eye field, LGN – lateral geniculate nucleus, M1 – primary motor cortex (M1).

## 5. Discussion

BPI allows us to simultaneously find ‘(de)activated’, ‘not activated’, and ‘low confidence’ voxels using a single decision rule. The ‘not activated’ decision means that the effect is negligible to an extent that it can be considered equivalent to the null for practical purposes. The ‘low confidence’ decision means we need more data to make a confident inference, that is, we need to increase the scanning time, sample size, or revisit the task design. The use of hierarchical PEB with the ‘global shrinkage’ prior enables us to check the results as the sample size increases and allows us to decide whether to optionally terminate the experiment if the obtained data are sufficient to make a confident inference. All the above features are absent from the classical NHST framework.

An important advantage of Bayesian inference is that we can use graphs such as those shown in Figure 12 to determine when the obtained data are sufficient to make a confident inference. We can plot such graphs for the whole brain or for a priori defined ROIs. When the curves reach a plateau, the data collection can be stopped. If the brain area can be labelled as either ‘(de)activated’ or ‘not activated’ at a relatively small sample size, it will be still so at larger sample sizes. If the brain area can be labelled as ‘low confidence’, we must increase the sample size to make a confident inference. At a certain sample size, it could possibly be labelled as either ‘(de)activated’ or ‘not activated’. In the worst case, we can reach the plateau and still label the brain area as ‘low confidence’. However, even in this case, we can make a definite conclusion: the task design is not sensitive to the effect and should be revised. Bayesian inference allows us to monitor the evidence for the alternative or null hypotheses after each participant without special adjustment for multiplicity (Edwards et al., 1963; Berger and Berry, 1988; Wagenmakers, 2007, Schönbrodt et al., 2015; Kruschke and Liddell, 2017b). The optional stopping of the experiment not only allows more freedom in terms of the experimental design, but also saves limited resources and is even more ethically justified in certain cases^3^ (Edwards et al., 1963; Wagenmakers, 2007).

In contrast, frequentist inference depends on the researcher’s intention to stop data collection and thus requires a definition of the stopping rule based on a priori power analysis. The sequential analysis and optional stopping in frequentist inference inflate the number of false positives. Moreover, even if the a priori defined sample size is reached, the researcher can still obtain a non-significant result. In this case, the researcher can follow two controversial paths within the classical NHST framework. Firstly, the sample size could be further increased to force an indecisive result to a decisive conclusion. The problem is that this conclusion would always be against the null hypothesis. Thus, an unbounded increase in the sample size introduces a discrepancy between classical NHST and Bayesian inference, also known as the Jeffreys-Lindley paradox. Secondly, one may argue that high a priori power and non-significant results provide evidence for the null hypothesis (see, for example, Cohen, 1990). However, even high *a priori* power and non-significant results do not provide direct evidence for the null hypothesis. In fact, a high-powered non-significant result may arise when the obtained data provide no evidence for the null over the alternative hypothesis, according to Bayesian inference (Denies and Mclatchie, 2017). This does not mean that power analysis is irrelevant from a Bayesian perspective. Although power analysis is not necessary for Bayesian inference, it can still be used within the Bayesian framework for study planning (Kruschke and Liddell, 2017b). At the same time, power analysis is a critical part of frequentist inference, as it depends on researcher intentions, such as the stopping intention.

The main difficulty with the application of BPI is the need to define the ES threshold. However, the problem of choosing a practically meaningful effect size is not unique to fMRI studies, as it arises in every mature field of science. It should not discourage us from using BPI, as the point-null hypothesis is never true in the soft sciences. From our perspective, there are several ways to address this problem. Firstly, the ES threshold can be chosen based on previously reported effect sizes in studies with a similar design or perform a pilot study to estimate the expected effect size.

Based on the fMRI literature, the largest BOLD responses are evoked by sensory stimulation and vary within 1–5% of the overall mean whole-brain activity. In contrast, BOLD responses induced by cognitive tasks vary within 0.1–0.5% (Friston et al., 2002b; Poldrack et al., 2011; Chen et al., 2017). The results obtained in this study support this notion. Primary sensory effects were >1%, and motor effects were >0.3%. Cognitive effects can be classified into three categories.

1. ‘Strong’ effects of 0.2–0.3% (for example, emotion processing in the amygdala, language processing in Broca’s area),
2. ‘Moderate’ effects of 0.1–0.2% (for example, working memory load in DLPFC, social cognition in IPL, response inhibition in IFG/FO),
3. ‘Weak’ effects of 0.05% in contrasts without robust activations (for example, reward processing in the nucleus accumbens, relational processing in DLPFC).

However, choosing the ES threshold based on relatively older studies can be challenging because fMRI designs become increasingly complex over time, and it can be difficult to find previous experiments reporting unbiased effect size with a similar design. In this case, one can use the ES threshold equal to *one prior SD* of the effect (Friston and Penny, 2003), which can be thought as a neuronal ‘background noise level’ or a level of activity that is generic to the whole brain (Eickhoff et al., 2008). BPI with this ES threshold generally works well for both ‘(de)activated’ and ‘not activated’ voxel detection. However, it may not be suitable in cases with very small differences between the dependent samples. In addition, researchers who rely more on the frequentist inference may use the *γ*(*Dice_max_*) threshold to replicate the results obtained previously with classical NHST and additionally search for ‘not activated’ and ‘low confidence’ voxels. Finally, the degree to which the posterior probability is contained within the ROPEs of different widths could be specified or the ROPE maps in which the thresholding sequence is inverted could be calculated. The ROPE maps can be shared in public repositories, such as Neurovault, along with PPMs, and subsequently thresholded by any reasonable ES threshold.

Classical NHST is limited to the detection of ‘(de)activated’ voxels. However, in a properly designed and conducted study, the ‘null effect’ can be just as valuable and exciting as rejecting the null hypothesis. Although the interval-based frequentist methods can be used to assess the ‘null effects’, they are less intuitive and more complicated in practice. At the same time, the results obtained with BPI have intuitively simple interpretations, as they provide direct evidence *for* the null or the alternative hypothesis, given the obtained data. Moreover, BPI has already been implemented in SPM12, which greatly facilitates its use by fMRI practitioners.

The ability to provide evidence for the null hypothesis may be especially beneficial for clinical neuroimaging. Possible issues that can be resolved using this approach are:

1. Let the brain activity in certain ROIs due to a neurodegenerative process decrease by more than *γ* per year on average without any treatment. To prove that a new treatment *effectively protects against neurodegenerative processes,* we can provide evidence that, within one year of treatment, brain activity was reduced by less than X%.
2. Assume that an effective treatment should change the brain activity in certain ROIs by at least X%. Then, we can prove that a new treatment is *practically ineffective* if the activity has changed by less than X%.
3. Consider two groups of subjects taking a new treatment and a placebo, respectively. Using BPI, we can provide evidence that the result of the new treatment is *does not differ from that of the placebo*.
4. Consider two groups of subjects taking an old effective treatment and a new treatment. Using BPI, we can provide evidence that the new treatment is *no worse than the old effective treatment*.
5. Consider a new treatment for a disease that *is not related to brain function*. Using BPI, we can provide evidence that the new treatment *does not have side effects* on brain activity.

## 6. Conclusions

Herein, a discussion of the use of the Bayesian and frequentist approaches to assess the ‘null effects’ was presented. We demonstrated that Bayesian inference may be more intuitive and convenient in practice than frequentist inference. Crucially, Bayesian inference can detect ‘(de)activated’, ‘not activated,’ and ‘low confidence’ voxels using a single decision rule. Moreover, it allows for interim analysis and optional stopping when the obtained sample size is sufficient to make a confident inference. We considered the problem of defining a threshold for the effect size and provided a reference set of typical effect sizes in different fMRI designs. Bayesian inference and assessment of the ‘null effects’ may be especially beneficial for basic and applied clinical neuroimaging. The developed SPM12-based scripts with a simple GUI is expected to be useful for the assessment of ‘null effects’ using BPI.

## 7. Limitations and future work

Firstly, we did not consider BMI, which is currently mainly used for the analysis of effective connectivity. A promising area of future research would be to compare the advantages of BMI and BPI when analysing local brain activity. Secondly, the ‘global shrinkage’ prior must be compared with other possible priors. Thirdly, we used Bayesian statistics only at the group level. Future studies could consider the advantages of using the Bayesian approach at both the subject and group levels.

## 8. Acknowledgments

Data were provided [in part] by the Human Connectome Project, WU-Minn Consortium (Principal Investigators: David Van Essen and Kamil Ugurbil; 1U54MH091657) funded by the 16 NIH Institutes and Centers that support the NIH Blueprint for Neuroscience Research; and by the McDonnell Center for Systems Neuroscience at Washington University. Another part of the data were provided by UCLA dataset which was obtained from the OpenfMRI database (its accession number is ds000030) and data collection was funded by the Consortium for Neuropsychiatric Phenomics (NIH Roadmap for Medical Research grants UL1-DE019580, RL1MH083268, RL1MH083269, RL1DA024853, RL1MH083270, RL1LM009833, PL1MH083271, and PL1NS062410). We thank Dr. Irina Knyazeva for her helpful discussions and valuable suggestions for this manuscript. We also thank Andrey Ogai for the valuable help with script for visualisation of statistical maps. RM, AK and MV were supported by the Russian Science Foundation grant #19-18-00454. YN, MD and DC were supported by the state assignment of the Ministry of Education and Science of Russian Federation (theme number AAAA-A19-119101890066-2).

## Footnotes

1 Here are some examples of ‘no effect’ conclusions that can be found in the fMRI literature: a) brain area was not activated, b) brain area was not involved in the function, c) no effect was found in the brain area (p>0.05), d) both groups showed no differences, which can be interpreted as evidence against the alternative hypothesis; e) patients have similar responses to both conditions (p>0.05), that is, they have difficulties in differentiating these conditions; f) lack of significant correlation during treatment suggest a protective impact of the therapy on brain areas.

2 FDR correction controls the rate of false discoveries (false positives in frequentist terminology) among all significant voxels. FWE correction controls the rate of any false positives in the whole brain.

3 This is especially true for PET studies. The BPI method described in this work can also be applied to PET data to reduce the sample size and thus exposure to radioactivity (Svensson et al., 2020).

