## Supplementary for "Evidence for the null hypothesis in functional magnetic resonance imaging using group-level Bayesian inference"

**Manuscript Title:**

### THEORY

Bayesian solution of the ‘Congressman example’ was proposed by Szucs and Ioannidis (2017).

1) The person citizenship is unknown a priori. That is, the prior probabilities of the null and alternative hypothesis are:

*P*(*H*_0_) = *P*(*H*_1_) = 0.5

2) As only American citizens can be a member of Congress, the conditional probability of the data (‘The person is a member of Congress’) under the alternative hypothesis is:

*P*(*D*|*H*_1_) = 0

3) There are 435 Congressmen out of 328 million American citizens. It is unlikely to be a member of Congress:

*P*(*D*|*H*_0_) = 10^-6^

4) Therefore, the posterior probability of the null hypothesis is:

$P\left( H_{0} | D \right)=\frac{P(H_{0})P\left( D | H_{0} \right)}{P\left( H_{0} \right)P\left( D | H_{0} \right)+P\left( H_{1} \right)P\left( D | H_{1} \right)}=\frac{0.5\cdot{10}^{-6}}{0.5\cdot{10}^{-6}+0.5\cdot0}=1$.

Using Bayesian inference we can accept the null hypothesis and conclude that the person is an American.

### METHODS

### 1. The fMRI data acquisition parameters

The fMRI data from the HCP dataset were collected using an echo-planar imaging (EPI) sequence on a modified 3T Siemens Skyra with TR (repetition time) = 720 ms, TE (echo time) = 33.1 ms, flip angle = 52°, in-plane FOV (field of view) = 208 × 180 mm, 72 slices, and voxel size = 2 × 2 × 2 mm, with a multi-band acceleration factor of 8 (Glasser et al., 2013). Each task consisted of two sessions, one with right-to-left and the other with left-to-right phase encoding. The fMRI data from the UCLA dataset were collected using an EPI sequence on a 3T Siemens Trio scanner with TR = 2 s, TE = 30 ms, flip angle = 90°, in-plane FOV = 192 × 192 mm, 34 slices, slice thickness = 4 mm, and voxel size = 3 × 3 × 4 mm (Poldrack et al., 2016). Each task consisted of one session.

### 2. Description of the general linear models used for each task design at the subject level of analysis

In the *emotion processing task*, each session consisted of three blocks with a presentation of emotional faces (fearful/angry) and three blocks in which the shapes were presented. Each block lasted 21 s. For each condition within a session, two regressors were included in a subject-level model. Each session included 10 seconds of fixation at the beginning.

In the *working memory (WM) task*, each session consisted of eight blocks for two WM load (0-back and 2-back) and four stimulus categories (faces, places, tools, body parts). Each block lasted 27.5 s. For each condition within a session, eight regressors were included in a subject-level model. Each session contained four fixation blocks (15 s).

In the *language task*, each session consisted of four blocks for the story condition interleaved with blocks for the math condition. On average, each block lasted for 30 s. For each condition within a session, two regressors were included in a subject-level model.

In the *motor task*, each session consisted of two blocks for tongue movement, two blocks for right/left hand movement, and two blocks for right/left foot movement. Each block lasted 12 s and preceded by a 3 s cue. For each condition within a session, ten regressors were included in a subject-level model (blocks and cues were modelled separately). Each session contained three fixation blocks (15 s).

In the *gambling task*, each session consisted of two blocks for mostly reward and two blocks for mostly loss conditions. Each block lasted 28 s. For each condition within a session, two regressors were included in a subject-level model. Each session contained four fixation blocks (15 s).

In the *social cognition task*, each session consisted of two or three blocks for the random interaction condition and two or three blocks for the mental interaction condition. Each block lasted 20 s. For each condition within a session, two regressors were included in a subject-level model. Each session contained five fixation blocks (15 s).

In the *relational processing task*, each session consisted of three blocks for the relational condition and three blocks for the matching condition. Each block lasted 16s. For each condition within a session, two regressors were included in a subject-level model. Each session contained three fixation blocks (16 s).

The *stop-signal task* (SST) consisted of 96 Go trials and 32 Stop-signal trials (approximately half of them were correct). Three regressors were included in a subject-level model with a fixed duration of 1.5 s. Jittered null events separated every trial.

The *task-switching task* consisted of 36 “Congruent, No Switch” trials, 12 “Congruent, Switch” trials, 36 “Incongruent, No Switch” trials, and 12 “Incongruent, Switch” trials. Four regressors were included in a subject-level model with a fixed duration of 1 s.

Summary information on the task considered in the current study presented in Table S1.

### Table S1. Summary information on the tasks considered in the current study.

| Task | Length (min:s) | Scans | Conditions | Contrasts | ROIs |
| --- | --- | --- | --- | --- | --- |
| **The HCP dataset, block designs (N = 100 subjects)** | | | | | |
| Emotion processing | 2:16 x 2 | 176 x 2 | Emotional faces  Shapes | Emotion > Shape | L/R Amy,  L/R Fus |
| Working memory | 5:01 x 2 | 405 x 2 | 0/2-back (faces, places, tools, back body parts)  Fixation | 2-back > baseline  2-back > 0-back | WM: L/R DLPFC,  L/R SPL, L/R IPL, dACC |
| Language  (auditory) | 3:57 x 2 | 316 x 2 | Story  Math | Story > Math | L IFG pt (BA45),  L IFG po (BA44),  L STG (Wernicke) |
| Motor | 3:34 x 2 | 284 x 2 | L/R finger (LF/RF)  L/R toe (LT/RT)  Tongue (T)  Fixation | LF > baseline  RF > baseline  T > baseline | L/R Precentral,  L/R Postcental,  SMA, L/R Put,  L/R Cerebellum |
| Gambling | 3:12 x 2 | 253 x 2 | Mostly reward  Mostly loss  Fixation | Reward > baseline  Loss > baseline  Reward > Loss | L/R NAc, L/R Amy, L/R Ins |
| Social cognition | 3:27 x 2 | 274 x 2 | Random interaction  Social interaction  Fixation | Social > baseline  Social > Random | L/R IPL, L/R STG,  L/R MTG |
| Relational  processing | 2:26 x 2 | 232 x 2 | Relational  Match  Fixation | Relational > baseline  Relational > Match | L/R DLPFC,  L/R SPL, L/R AI/FO,  preSMA/dACC |
| **The UCLA dataset, event-related designs (N = 115 subjects)** | | | | | |
| Stop-Signal task (auditory stop-signal) | 6:08 | 184 | Go  Stop-signal (~50/50 – correct/incorrect)  Null events | Correct Stop > baseline  Correct Stop > Go | L/R IFG/FO,  L/R DLPFC,  preSMA/dACC |
| Task-switching | 6:56 | 208 | Congruent, no switch  Congruent, switch  Incongruent, no switch  Incongruent, switch | Switch > baseline  Switch > No switch | L/R DLPFC,  L/R SPL, L/R LOC,  preSMA/dACC,  L/R AI/FO |

*L/R – left/right, Amy – amygdala, Fus – fusiform gyrus, DLPFC – dorsolateral prefrontal cortex, SPL – superior parietal lobule, IPL – inferior parietal lobule, dACC – dorsal anterior cingulate cortex, IFG – inferior frontal gyrus, pt – pars triangularis, po – pars opercularis, BA – Brodmann area, STG – superior temporal gyrus, SMA – supplementary motor area, put – putamen, NAc – nucleus accumbens, Ins – insular cortex, MTG – middle temporal gyrus, AI – anterior insula, FO – frontal operculum, LOC – lateral occipital cortex.*

### 3. Calculation of BOLD percent signal change

Raw beta values calculated by SPM12 at the subject level of analysis represent the BOLD signal in arbitrary units. They can be scaled to the BOLD percent signal change (PSC) relative to the mean whole-brain “baseline” signal. In the current work, BOLD PSC was calculated following the procedure recommended in Poldrack, Mumford and Nichols (2011, p.186):

$${PSC}_{i}=SF*\frac{{\hat{\beta}cond}_{i}}{\sum_{i=1}^{n} {\hat{\beta}const}_{i}/ n}*100\%$$

${\hat{\beta}cond}_{i}$is the parameter estimate for *i^th^* voxel for a certain condition.

${\hat{\beta}const}_{i}$is the parameter estimate for *i^th^* voxel for the constant term in a certain task session. It represents the mean BOLD signal for all time points within the session, adjusted for all other effects modelled in the subject level design matrix (all experimental conditions and nuisance regressors).

The denominator represents ${\hat{\beta}const}_{i}$ averaged over all *n* voxels within the brain. In SPM12 the constant term can be interpreted as “implicit baseline” or the activity in the “resting state condition” (fixation) which are not modeled explicitly (Pernet, 2014).

SF is the scaling factor applied for making the peak of an isolated BOLD response equal to one. For the task design with event duration of 0 s, the SF can be calculated using information stored in “SPM.mat”: SF = max(SPM.xBF.bf(:,1))/SPM.xBF.dt. To find the peak of isolated BOLD response in other cases, we can create the upsampled regressor in a higher time resolution for a single event with a specified duration. It can be done via ‘fMRI model specification (design only)’ module in SPM batch editor.

The SF for all block-design tasks was 1.1443, the SF for the stop-signal task was 0.3212, and the SF for the task-switching task was 0.2113.

The linear contrast of beta values scaled to PSC should have a sum of positive terms equals to 1 and a sum of negative terms equals to -1.

### 4. Creation of ROI masks

A priori ROI masks were used to estimate unbiased typical effect sizes in different task designs. ROI masks were defined as the overlap between anatomical and a priori functional masks (Poldrack et al., 2017). Functional masks were obtained using Neurosynth platform for an automatic meta-analysis (Yarkoni et al., 2011). A uniformity test with default false-discovery rate (FDR) corrected p<0.01 threshold was chosen. For the Neurosynth meta-analysis, the following search terms were used:

1) Emotion processing task – ‘emotion’ (1037 studies) and ‘face’ (896 studies);
2) Working memory task – ‘working memory’ (1091 studies);
3) Language task – ‘language’ (1101 studies);
4) Motor task – ‘motor’ (2565 studies);
5) Gambling task – ‘reward’ (922 studies);
6) Social cognition – ‘social’ (1302 studies);
7) Relational processing – ‘relational’ (91 studies);
8) Stop-signal task – ‘response inhibition’ (218 studies);
9) Task-switching – ‘switching’ (193 studies);
10) Additionally, for tasks with visual stimuli – ‘visual’ (3110 studies).
11) Additionally, for tasks with auditory stimuli – ‘auditory’ (1252 studies).

To obtain anatomical masks, we used discrete labels from the Harvard-Oxford atlas (Desikan et al., 2006) derived from taking the structure with the maximum probability at each location. In comparison, Poldrack et al. (2017) used the Harvard–Oxford probabilistic atlas at P > 0, which results in much larger ROIs overlapping with several adjacent anatomical brain regions. For example, the ‘right amygdala ROI’ consisted of 1082 voxels, i.e. ROI volume was equal 8656 mm^3^ (voxel size 2 mm^3^) or 8.6 cm^3^ which is too large for the amygdala. In healthy adults, right amygdala volume reaches 1.88 cm^3^ (Rice et al., 2014). ROIs obtained by P > 0 threshold span over multiple regions which reduces neurophysiological interpretability of effect sizes extracted from them. The selection of anatomical masks for each task was based on the previously reported results specific to a certain type of tasks. The anatomical masks with the following labels from the Harvard-Oxford atlas were used:

1) Amygdala – ‘amygdala’;
2) Fusiform gyrus – ‘occipital fusiform gyrus’ and ‘temporal fusiform gyrus’;
3) Dorsolateral prefrontal cortex – ‘middle frontal gyrus’ and ‘frontal pole’;
4) Superior parietal lobule – ‘superior parietal lobule’;
5) Inferior parietal lobule – ‘supramarginal anterior’, ‘supramarginal posterior’, and ‘angular gyrus’;
6) Dorsal anterior cingulate cortex – ‘paracingulate gyrus’;
7) Inferior frontal gyrus, pars triangularis – ‘inferior frontal gyrus, pars triangularis’;
8) Inferior frontal gyrus, pars opercularis – ‘inferior frontal gyrus, pars opercularis’;
9) Superior temporal gyrus – ‘superior temporal gyrus, posterior division’ and ‘planum temporale’;
10) Precentral gyrus – ‘precentral gyrus’;
11) Postcental gyrus – ‘precentral gyrus’;
12) Supplementary motor area – ‘juxtapositional lobule cortex’;
13) Putamen – ‘putamen’;
14) Nucleus accumbens – ‘accumbens’;
15) Insular cortex – ‘insular cortex’;
16) Middle temporal gyrus – ‘Middle Temporal Gyrus, anterior division’, ‘posterior division’, and ‘temporooccipital part’;
17) Anterior insula, frontal operculum (AIFO) – ‘frontal operculum cortex’ and ‘insular cortex’;
18) Lateral occipital cortex – ‘lateral occipital cortex, superior division’.

To obtain anatomical masks of the cerebellum WFU Pickatlas toolbox was used (Maldjian et al., 2003). Additionally, for each task with visual stimuli, the effect sizes from primary and secondary visual areas (V1 and V2) ware reported. V1 and V2 ROIs were obtained using the SPM Anatomy toolbox (Eickhoff et al., 2005). For tasks with auditory stimuli, the effect sizes from the primary auditory area (Heschl's gyri from the Harvard-Oxford atlas) were reported additionally. See the list of selected ROIs for each task in Table S1.

### RESULTS

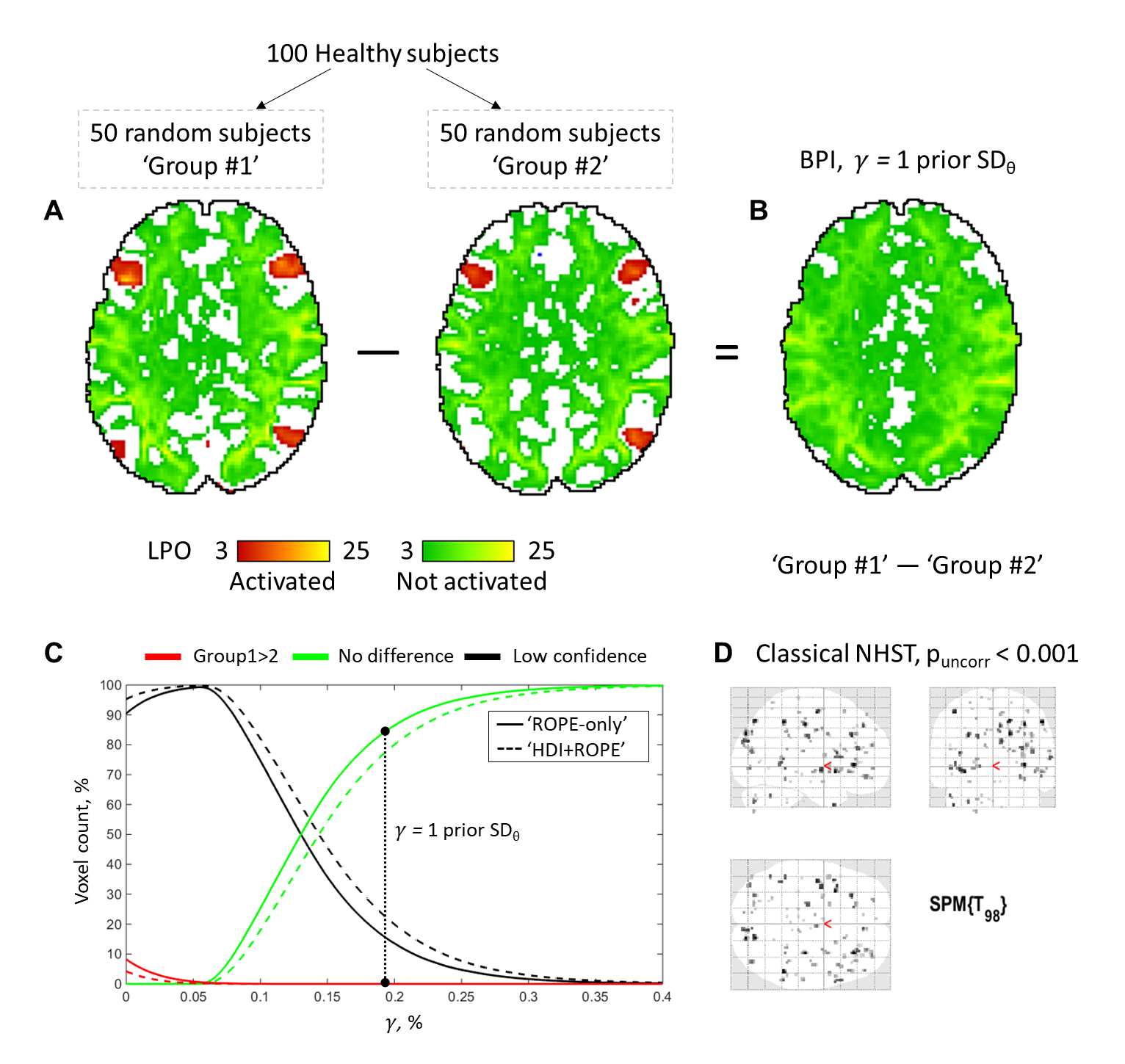

### Figure S1. Comparison of two independent samples with ‘similar’ levels of neuronal activity.

To emulate two ‘similar’ *independent* samples, 100 healthy adult subjects’ contrasts (‘Emotion > Shape’) were randomly assigned into two groups of 50 subjects. A) Within-group results obtained using BPI with the ‘ROPE-only’ decision rule and *γ = 1* *prior SD_θ_* threshold. Axial slice z = 20 mm (MNI152 standard space). B) Results of the comparison of two independent samples obtained using BPI with the ‘ROPE-only’ decision rule and *γ = 1* *prior SD_θ_* threshold. C) Dependence between the ES threshold γ and the amount of ‘no difference’ and ‘low confidence’ voxels when comparing two independent samples using the ‘ROPE-only’ rule (solid lines) and ‘HDI+ROPE’ rule (dashed lines). D) Results of the comparison of two independent samples using classical NHST. An uncorrected p < 0.001 threshold was used for illustrational purposes.

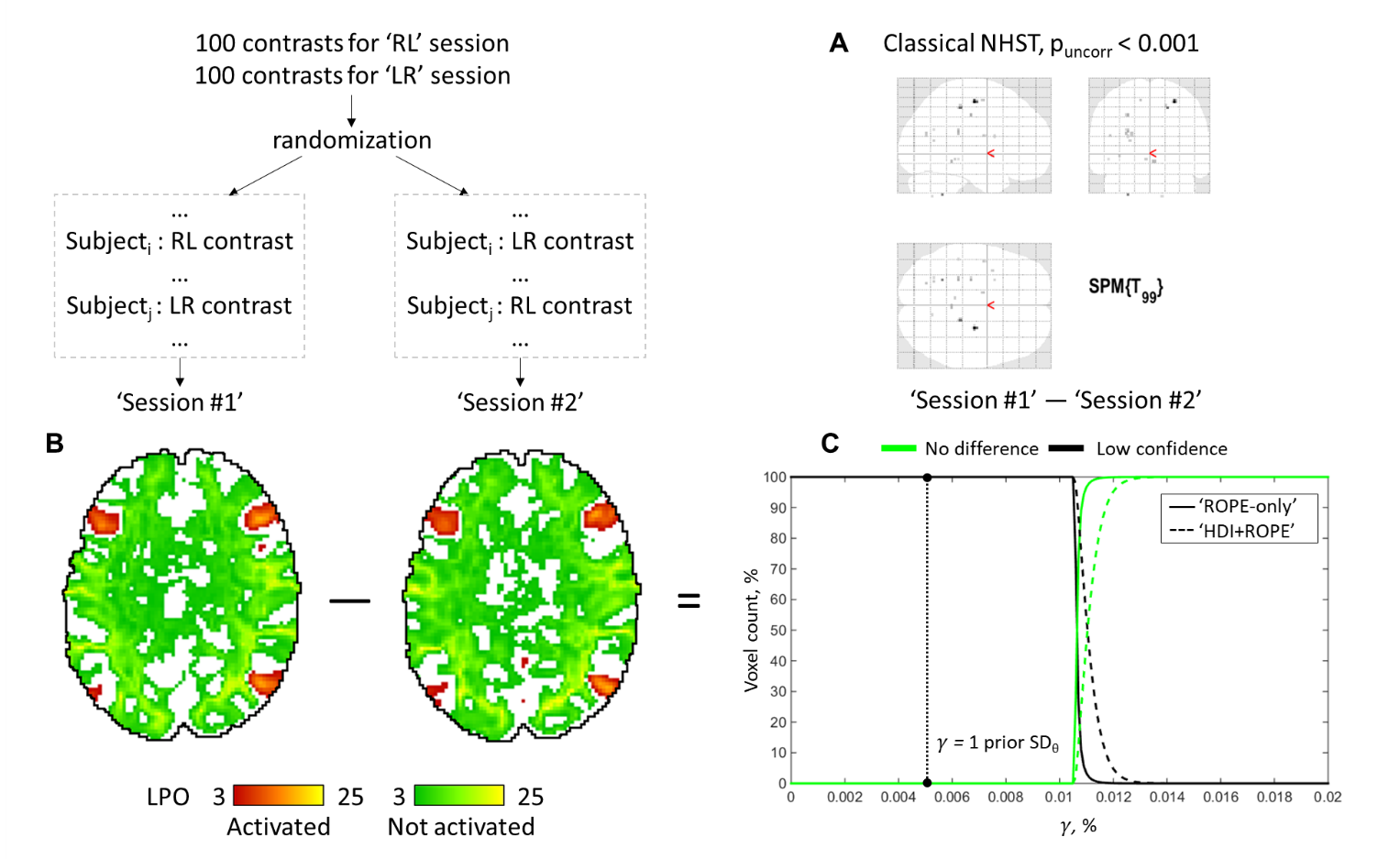

### Figure S2. Comparison of two dependent samples with ‘similar’ levels of neuronal activity.

To emulate ‘similar’ *dependent* samples, we randomized ‘Emotion > Shape’ contrasts from right-to-left (RL) and left-to-right (LR) phase encoding sessions in the ‘Session #1’ and ‘Session #2’ samples. Each sample consisted of 50 contrasts from the RL session and 50 from the LR session. A) Results of the comparison of two dependent samples using classical NHST. An uncorrected p < 0.001 threshold was used for illustrational purposes. B) Within-session results obtained using BPI with the ‘ROPE-only’ decision rule and *γ = 1* *prior SD_θ_* threshold. Axial slice z = 20 mm (MNI152 standard space). C) Dependence between the ES threshold γ and the amount of ‘no difference’ and ‘low confidence’ voxels when comparing two dependent samples using the ‘ROPE-only’ rule (solid lines) and ‘HDI+ROPE’ rule (dashed lines).

### Table S2. The emotion processing task results.

Comparison of classical NHST with voxel-wise FWE correction (p<0.05) and Bayesian parameter inference with different ES thresholds. Amount of voxels in % from the total number of voxels.

| Classification | Classical  NHST | *γ(Dice_max_)* | *γ = 0.1%* | *γ = 0.2*  *prior SD_θ_* | *γ = 0.5*  *prior SD_θ_* | *γ = 0.8*  *prior SD_θ_* | *γ = 1*  *prior SD_θ_* |
| --- | --- | --- | --- | --- | --- | --- | --- |
| **Emotion > Shape. The ‘ROPE-only’ decision rule. *1 prior SD_θ_ = 0.135%*** | | | | | | | |
| Activation | 8.70 | 8.73 | 10.13 | 22.12 | 13.90 | 9.41 | 7.40 |
| Deactivation | 0.54 | 0.12 | 0.26 | 7.16 | 1.28 | 0.19 | 0.04 |
| No activation | - | 67.54 | 58.82 | 0.12 | 32.38 | 63.45 | 75.03 |
| Low-confidence | - | 23.62 | 30.79 | 70.60 | 52.45 | 26.96 | 17.52 |
| **Emotion > Shape. The ‘HDI+ROPE’ decision rule. *1 prior SD_θ_ = 0.135%*** | | | | | | | |
| Activation | 8.70 | 8.80 | 9.14 | 19.71 | 12.56 | 8.51 | 6.72 |
| Deactivation | 0.54 | 0.14 | 0.17 | 5.38 | 0.80 | 0.11 | 0.02 |
| No activation | - | 55.70 | 53.05 | 0.05 | 26.21 | 58.13 | 71.39 |
| Low-confidence | - | 35.37 | 37.64 | 74.86 | 60.42 | 33.24 | 21.87 |

### Table S3. The working memory task results.

Comparison of classical NHST with voxel-wise FWE correction (p<0.05) and Bayesian parameter inference with different ES thresholds. Amount of voxels in % from the total number of voxels.

| Classification | Classical  NHST | *γ(Dice_max_)* | *γ = 0.1%* | *γ = 0.2*  *prior SD_θ_* | *γ = 0.5*  *prior SD_θ_* | *γ = 0.8*  *prior SD_θ_* | *γ = 1*  *prior SD_θ_* |
| --- | --- | --- | --- | --- | --- | --- | --- |
| **2-back > baseline. The ‘ROPE-only’ decision rule. *1 prior SD_θ_ = 0.325%*** | | | | | | | |
| Activation | 26.23 | 27.04 | 31.03 | 35.72 | 24.49 | 17.25 | 13.79 |
| Deactivation | 9.70 | 8.51 | 11.39 | 15.03 | 6.92 | 2.93 | 1.72 |
| No activation | - | 36.25 | 24.06 | 9.77 | 43.59 | 63.23 | 71.89 |
| Low-confidence | - | 28.21 | 33.52 | 39.48 | 25.00 | 16.59 | 12.60 |
| **2-back > baseline. The ‘HDI+ROPE’ decision rule. *1 prior SD_θ_ = 0.325%*** | | | | | | | |
| Activation | 26.23 | 26.84 | 29.56 | 33.99 | 23.30 | 16.38 | 13.09 |
| Deactivation | 9.70 | 8.51 | 10.45 | 13.87 | 6.30 | 2.62 | 1.54 |
| No activation | - | 29.34 | 20.72 | 7.24 | 40.56 | 61.15 | 70.31 |
| Low-confidence | - | 35.31 | 39.27 | 44.90 | 29.84 | 19.85 | 15.06 |
| **2-back > 0-back. The ‘ROPE-only’ decision rule. *1 prior SD_θ_ = 0.089%*** | | | | | | | |
| Activation | 6.56 | 6.75 | 6.29 | 17.71 | 12.60 | 9.10 | 7.25 |
| Deactivation | 4.17 | 3.28 | 2.86 | 19.56 | 11.39 | 6.12 | 3.82 |
| No activation | - | 52.69 | 56.00 | 0.00 | 8.46 | 34.34 | 48.98 |
| Low-confidence | - | 37.29 | 34.85 | 62.74 | 67.56 | 50.44 | 39.95 |
| **2-back > 0-back. The ‘HDI+ROPE’ decision rule. *1 prior SD_θ_ = 0.089%*** | | | | | | | |
| Activation | 6.56 | 6.42 | 5.51 | 15.71 | 11.21 | 8.08 | 6.37 |
| Deactivation | 4.17 | 3.02 | 2.22 | 16.79 | 9.53 | 4.90 | 2.98 |
| No activation | - | 42.47 | 50.34 | 0.00 | 5.20 | 28.08 | 42.95 |
| Low-confidence | - | 48.08 | 41.93 | 67.50 | 74.06 | 58.93 | 47.71 |

### Table S4. The language task results.

Comparison of classical NHST with voxel-wise FWE correction (p<0.05) and Bayesian parameter inference with different ES thresholds. Amount of voxels in % from the total number of voxels.

| Classification | Classical  NHST | *γ(Dice_max_)* | *γ = 0.1%* | *γ = 0.2*  *prior SD_θ_* | *γ = 0.5*  *prior SD_θ_* | *γ = 0.8*  *prior SD_θ_* | *γ = 1*  *prior SD_θ_* |
| --- | --- | --- | --- | --- | --- | --- | --- |
| **Story > Math. The ‘ROPE-only’ decision rule. *1 prior SD_θ_ = 0.255%*** | | | | | | | |
| Activation | 12.6 | 13.2 | 14.8 | 20.2 | 12.6 | 8.5 | 6.6 |
| Deactivation | 16.9 | 19.1 | 21.4 | 29.5 | 18.1 | 12.2 | 9.6 |
| No activation | - | 31.3 | 23.4 | 3.9 | 34.7 | 57.8 | 67.8 |
| Low-confidence | - | 36.4 | 40.4 | 46.4 | 34.5 | 21.5 | 16.0 |
| **Story > Math. The ‘HDI+ROPE’ decision rule. *1 prior SD_θ_ = 0.255%*** | | | | | | | |
| Activation | 12.6 | 13.2 | 13.8 | 18.9 | 11.8 | 7.9 | 6.2 |
| Deactivation | 16.9 | 18.9 | 19.8 | 27.3 | 16.8 | 11.3 | 8.9 |
| No activation | - | 22.9 | 19.7 | 2.5 | 30.6 | 54.8 | 65.4 |
| Low-confidence | - | 45.1 | 46.7 | 51.3 | 40.7 | 25.9 | 19.4 |

### Table S5. The motor task results.

Comparison of classical NHST with voxel-wise FWE correction (p<0.05) and Bayesian parameter inference with different ES thresholds. Amount of voxels in % from the total number of voxels.

| Classification | Classical  NHST | *γ(Dice_max_)* | *γ = 0.1%* | *γ = 0.2*  *prior SD_θ_* | *γ = 0.5*  *prior SD_θ_* | *γ = 0.8*  *prior SD_θ_* | *γ = 1*  *prior SD_θ_* |
| --- | --- | --- | --- | --- | --- | --- | --- |
| **Left finger > baseline. The ‘ROPE-only’ decision rule. *1 prior SD_θ_ = 0.149%*** | | | | | | | |
| Activation | 5.2 | 5.1 | 7.4 | 13.5 | 9.1 | 6.4 | 5.1 |
| Deactivation | 3.2 | 3.1 | 6.7 | 18.7 | 9.8 | 5.0 | 3.1 |
| No activation | - | 55.0 | 28.4 | 0.0 | 11.3 | 40.3 | 55.4 |
| Low-confidence | - | 36.7 | 57.5 | 67.8 | 69.8 | 48.4 | 36.3 |
| **Left finger > baseline. The ‘HDI+ROPE’ decision rule. *1 prior SD_θ_ = 0.149%*** | | | | | | | |
| Activation | 5.2 | 5.1 | 6.6 | 11.9 | 8.1 | 5.7 | 4.6 |
| Deactivation | 3.2 | 3.0 | 5.4 | 15.4 | 7.9 | 4.0 | 2.4 |
| No activation | - | 42.4 | 22.1 | 0.0 | 7.2 | 33.5 | 49.3 |
| Low-confidence | - | 49.5 | 65.9 | 72.7 | 76.7 | 56.7 | 43.7 |
| **Right finger > baseline. The ‘ROPE-only’ decision rule. *1 prior SD_θ_ = 0.171%*** | | | | | | | |
| Activation | 8.0 | 8.0 | 12.2 | 20.0 | 13.6 | 9.4 | 7.4 |
| Deactivation | 1.7 | 1.9 | 4.6 | 12.1 | 5.7 | 2.7 | 1.6 |
| No activation | - | 52.0 | 20.4 | 0.0 | 11.8 | 41.1 | 56.4 |
| Low-confidence | - | 38.1 | 62.8 | 67.9 | 68.9 | 46.8 | 34.6 |
| **Right finger > baseline. The ‘HDI+ROPE’ decision rule. *1 prior SD_θ_ = 0.171%*** | | | | | | | |
| Activation | 8.0 | 8.0 | 10.9 | 17.7 | 12.1 | 8.4 | 6.7 |
| Deactivation | 1.7 | 1.9 | 3.6 | 9.8 | 4.5 | 2.1 | 1.3 |
| No activation | - | 38.1 | 14.8 | 0.0 | 7.5 | 34.3 | 50.2 |
| Low-confidence | - | 52.0 | 70.7 | 72.5 | 75.9 | 55.1 | 41.8 |
| **Tongue > baseline. The ‘ROPE-only’ decision rule. *1 prior SD_θ_ = 0.268%*** | | | | | | | |
| Activation | 6.5 | 6.5 | 10.1 | 12.8 | 8.7 | 6.3 | 5.3 |
| Deactivation | 6.6 | 6.9 | 15.9 | 22.9 | 12.1 | 6.4 | 4.2 |
| No activation | - | 45.6 | 6.5 | 0.0 | 19.7 | 48.3 | 61.8 |
| Low-confidence | - | 41.0 | 67.5 | 64.3 | 59.5 | 38.9 | 28.8 |
| **Tongue > baseline. The ‘HDI+ROPE’ decision rule. *1 prior SD_θ_ = 0.268%*** | | | | | | | |
| Activation | 6.5 | 6.7 | 9.3 | 11.5 | 8.1 | 5.9 | 4.9 |
| Deactivation | 6.6 | 7.0 | 13.7 | 19.8 | 10.3 | 5.4 | 3.5 |
| No activation | - | 31.9 | 3.9 | 0.0 | 14.4 | 42.8 | 57.1 |
| Low-confidence | - | 54.4 | 73.1 | 68.6 | 67.3 | 46.0 | 34.5 |

### Table S6. The gabling task results.

Comparison of classical NHST with voxel-wise FWE correction (p<0.05) and Bayesian parameter inference with different ES thresholds. Amount of voxels in % from the total number of voxels.

| Classification | Classical  NHST | *γ(Dice_max_)* | *γ = 0.1%* | *γ = 0.2*  *prior SD_θ_* | *γ = 0.5*  *prior SD_θ_* | *γ = 0.8*  *prior SD_θ_* | *γ = 1*  *prior SD_θ_* |
| --- | --- | --- | --- | --- | --- | --- | --- |
| **Reward > baseline. The ‘ROPE-only’ decision rule. *1 prior SD_θ_ = 0.254%*** | | | | | | | |
| Activation | 24.0 | 24.2 | 28.3 | 36.2 | 24.7 | 17.3 | 13.9 |
| Deactivation | 7.4 | 6.3 | 8.7 | 14.2 | 6.7 | 2.8 | 1.5 |
| No activation | - | 39.8 | 27.8 | 5.1 | 38.2 | 59.4 | 68.9 |
| Low-confidence | - | 29.6 | 35.2 | 44.5 | 30.4 | 20.4 | 15.7 |
| **Reward > baseline. The ‘HDI+ROPE’ decision rule. *1 prior SD_θ_ = 0.254%*** | | | | | | | |
| Activation | 24.0 | 23.9 | 26.7 | 34.2 | 23.3 | 16.4 | 13.1 |
| Deactivation | 7.4 | 6.3 | 7.9 | 12.9 | 5.9 | 2.4 | 1.2 |
| No activation | - | 32.7 | 24.0 | 3.3 | 34.6 | 56.8 | 66.9 |
| Low-confidence | - | 37.1 | 41.4 | 49.6 | 36.2 | 24.3 | 18.8 |
| **Loss > baseline. The ‘ROPE-only’ decision rule. *1 prior SD_θ_ = 0.249%*** | | | | | | | |
| Activation | 20.1 | 20.1 | 23.8 | 31.0 | 21.0 | 14.9 | 12.1 |
| Deactivation | 10.7 | 9.2 | 12.5 | 19.3 | 10.0 | 5.1 | 3.1 |
| No activation | - | 40.0 | 26.9 | 4.3 | 36.6 | 58.6 | 68.5 |
| Low-confidence | - | 30.7 | 36.8 | 45.4 | 32.3 | 21.4 | 16.3 |
| **Loss > baseline. The ‘HDI+ROPE’ decision rule. *1 prior SD_θ_ = 0.249%*** | | | | | | | |
| Activation | 20.1 | 20.4 | 22.4 | 29.2 | 19.8 | 14.1 | 11.4 |
| Deactivation | 10.7 | 9.7 | 11.3 | 17.8 | 9.1 | 4.5 | 2.7 |
| No activation | - | 30.3 | 23.1 | 2.6 | 32.7 | 55.8 | 66.3 |
| Low-confidence | - | 39.5 | 43.1 | 50.4 | 38.4 | 25.6 | 19.7 |
| **Reward > Loss. The ‘ROPE-only’ decision rule. *1 prior SD_θ_ = 0.032%*** | | | | | | | |
| Activation | 0.5 | 0.5 | 0.0 | 7.3 | 3.3 | 1.5 | 1.0 |
| Deactivation | 0.0 | 0.0 | 0.0 | 0.1 | 0.0 | 0.0 | 0.0 |
| No activation | - | 11.8 | 98.1 | 0.0 | 0.0 | 0.0 | 0.0 |
| Low-confidence | - | 87.7 | 1.9 | 92.7 | 96.7 | 98.5 | 99.0 |
| **Reward > Loss. The ‘HDI+ROPE’ decision rule. *1 prior SD_θ_ = 0.032%*** | | | | | | | |
| Activation | 0.5 | 0.5 | 0.0 | 4.0 | 1.9 | 0.9 | 0.6 |
| Deactivation | 0.0 | 0.0 | 0.0 | 0.0 | 0.0 | 0.0 | 0.0 |
| No activation | - | 0.0 | 96.1 | 0.0 | 0.0 | 0.0 | 0.0 |
| Low-confidence | - | 99.5 | 3.9 | 95.9 | 98.1 | 99.1 | 99.4 |

### Table S7. The social cognition task results.

Comparison of classical NHST with voxel-wise FWE correction (p<0.05) and Bayesian parameter inference with different ES thresholds. Amount of voxels in % from the total number of voxels.

| Classification | Classical  NHST | *γ(Dice_max_)* | *γ = 0.1%* | *γ = 0.2*  *prior SD_θ_* | *γ = 0.5*  *prior SD_θ_* | *γ = 0.8*  *prior SD_θ_* | *γ = 1*  *prior SD_θ_* |
| --- | --- | --- | --- | --- | --- | --- | --- |
| **Social interaction > baseline. The ‘ROPE-only’ decision rule. *1 prior SD_θ_ = 0.325%*** | | | | | | | |
| Activation | 33.8 | 34.1 | 38.9 | 43.9 | 31.6 | 23.4 | 19.3 |
| Deactivation | 4.9 | 5.5 | 7.4 | 9.9 | 4.7 | 2.2 | 1.3 |
| No activation | - | 32.9 | 19.9 | 6.9 | 39.2 | 57.6 | 66.0 |
| Low-confidence | - | 27.5 | 33.8 | 39.3 | 24.5 | 16.8 | 13.5 |
| **Social interaction > baseline. The ‘HDI+ROPE’ decision rule. *1 prior SD_θ_ = 0.325%*** | | | | | | | |
| Activation | 33.8 | 34.4 | 37.3 | 42.1 | 30.3 | 22.5 | 18.4 |
| Deactivation | 4.9 | 5.6 | 6.6 | 8.8 | 4.2 | 1.9 | 1.1 |
| No activation | - | 24.9 | 16.6 | 4.9 | 36.1 | 55.5 | 64.3 |
| Low-confidence | - | 35.2 | 39.4 | 44.1 | 29.4 | 20.1 | 16.2 |
| **Social interaction > Random interaction. The ‘ROPE-only’ decision rule. *1 prior SD_θ_ = 0.104%*** | | | | | | | |
| Activation | 8.1 | 8.2 | 9.5 | 21.1 | 15.5 | 11.4 | 9.1 |
| Deactivation | 2.1 | 1.8 | 2.4 | 12.5 | 6.6 | 3.4 | 2.2 |
| No activation | - | 48.7 | 39.5 | 0.0 | 3.7 | 26.5 | 42.1 |
| Low-confidence | - | 41.3 | 48.6 | 66.4 | 74.2 | 58.8 | 46.6 |
| **Social interaction > Random interaction. The ‘HDI+ROPE’ decision rule. *1 prior SD_θ_ = 0.104%*** | | | | | | | |
| Activation | 8.1 | 8.0 | 8.4 | 18.8 | 13.9 | 10.1 | 8.0 |
| Deactivation | 2.1 | 1.8 | 1.9 | 9.9 | 5.2 | 2.7 | 1.8 |
| No activation | - | 35.2 | 32.5 | 0.0 | 1.8 | 20.1 | 35.1 |
| Low-confidence | - | 55.0 | 57.2 | 71.2 | 79.2 | 67.1 | 55.1 |

### Table S8. The relational processing task results.

Comparison of classical NHST with voxel-wise FWE correction (p<0.05) and Bayesian parameter inference with different ES thresholds. Amount of voxels in % from the total number of voxels.

| Classification | Classical  NHST | *γ(Dice_max_)* | *γ = 0.1%* | *γ = 0.2*  *prior SD_θ_* | *γ = 0.5*  *prior SD_θ_* | *γ = 0.8*  *prior SD_θ_* | *γ = 1*  *prior SD_θ_* |
| --- | --- | --- | --- | --- | --- | --- | --- |
| **Relational > baseline. The ‘ROPE-only’ decision rule. *1 prior SD_θ_ = 0.390%*** | | | | | | | |
| Activation | 28.7 | 29.0 | 34.4 | 37.0 | 25.6 | 18.3 | 14.9 |
| Deactivation | 8.9 | 8.0 | 12.3 | 14.7 | 5.7 | 2.2 | 1.2 |
| No activation | - | 34.6 | 19.0 | 11.6 | 44.5 | 64.4 | 73.0 |
| Low-confidence | - | 28.3 | 34.2 | 36.7 | 24.1 | 15.1 | 10.9 |
| **Relational > baseline. The ‘HDI+ROPE’ decision rule. *1 prior SD_θ_ = 0.390%*** | | | | | | | |
| Activation | 28.7 | 29.0 | 34.4 | 37.0 | 25.6 | 18.3 | 14.9 |
| Deactivation | 8.9 | 8.0 | 12.3 | 14.7 | 5.7 | 2.2 | 1.2 |
| No activation | - | 34.6 | 19.0 | 11.6 | 44.5 | 64.4 | 73.0 |
| Low-confidence | - | 28.3 | 34.2 | 36.7 | 24.1 | 15.1 | 10.9 |
| **Relational > Match. The ‘ROPE-only’ decision rule. *1 prior SD_θ_ = 0.051%*** | | | | | | | |
| Activation | 1.1 | 1.0 | 0.4 | 8.1 | 5.4 | 3.5 | 2.4 |
| Deactivation | 1.0 | 0.6 | 0.0 | 20.0 | 11.0 | 5.1 | 2.9 |
| No activation | - | 49.0 | 76.8 | 0.0 | 0.0 | 4.9 | 17.8 |
| Low-confidence | - | 49.4 | 22.9 | 71.9 | 83.6 | 86.5 | 76.9 |
| **Relational > Match. The ‘HDI+ROPE’ decision rule. *1 prior SD_θ_ = 0.051%*** | | | | | | | |
| Activation | 1.1 | 0.9 | 0.3 | 6.6 | 4.3 | 2.6 | 1.7 |
| Deactivation | 1.0 | 0.6 | 0.0 | 15.2 | 7.8 | 3.4 | 1.8 |
| No activation | - | 31.1 | 69.8 | 0.0 | 0.0 | 2.3 | 11.3 |
| Low-confidence | - | 67.5 | 29.9 | 78.2 | 87.9 | 91.7 | 85.2 |

### Table S9. The stop-signal task results.

Comparison of classical NHST with voxel-wise FWE correction (p<0.05) and Bayesian parameter inference with different ES thresholds. Amount of voxels in % from the total number of voxels.

| Classification | Classical  NHST | *γ(Dice_max_)* | *γ = 0.1%* | *γ = 0.2*  *prior SD_θ_* | *γ = 0.5*  *prior SD_θ_* | *γ = 0.8*  *prior SD_θ_* | *γ = 1*  *prior SD_θ_* |
| --- | --- | --- | --- | --- | --- | --- | --- |
| **Correct Stop > baseline. The ‘ROPE-only’ decision rule. *1 prior SD_θ_ = 0.069%*** | | | | | | | |
| Activation | 7.1 | 7.3 | 3.5 | 23.1 | 14.2 | 9.1 | 6.9 |
| Deactivation | 2.9 | 3.4 | 1.5 | 12.4 | 7.6 | 4.6 | 3.2 |
| No activation | - | 48.3 | 72.2 | 0.0 | 9.3 | 36.8 | 50.8 |
| Low-confidence | - | 41.0 | 22.9 | 64.4 | 69.0 | 49.6 | 39.0 |
| **Correct Stop > baseline. The ‘HDI+ROPE’ decision rule. *1 prior SD_θ_ = 0.069%*** | | | | | | | |
| Activation | 7.1 | 6.9 | 2.9 | 19.8 | 12.3 | 7.9 | 5.8 |
| Deactivation | 2.9 | 3.2 | 1.1 | 10.4 | 6.3 | 3.7 | 2.6 |
| No activation | - | 37.2 | 68.1 | 0.0 | 4.8 | 30.6 | 45.0 |
| Low-confidence | - | 52.7 | 27.8 | 69.8 | 76.6 | 57.9 | 46.6 |
| **Correct Stop > Go. The ‘ROPE-only’ decision rule. *1 prior SD_θ_ = 0.064%*** | | | | | | | |
| Activation | 12.8 | 13.2 | 5.2 | 28.0 | 19.2 | 13.5 | 10.4 |
| Deactivation | 2.5 | 2.6 | 0.6 | 11.2 | 5.3 | 2.7 | 1.8 |
| No activation | - | 48.8 | 80.0 | 0.0 | 20.7 | 47.7 | 59.6 |
| Low-confidence | - | 35.4 | 14.1 | 60.8 | 54.8 | 36.2 | 28.2 |
| **Correct Stop > Go. The ‘HDI+ROPE’ decision rule. *1 prior SD_θ_ = 0.064%*** | | | | | | | |
| Activation | 12.8 | 13.0 | 4.6 | 25.2 | 17.4 | 12.0 | 9.3 |
| Deactivation | 2.5 | 2.5 | 0.5 | 9.2 | 4.3 | 2.2 | 1.5 |
| No activation | - | 37.5 | 77.7 | 0.0 | 15.3 | 42.5 | 55.3 |
| Low-confidence | - | 47.0 | 17.2 | 65.6 | 63.0 | 43.2 | 33.9 |

### Table S10. The task-switching task results.

Comparison of classical NHST with voxel-wise FWE correction (p<0.05) and Bayesian parameter inference with different ES thresholds. Amount of voxels in % from the total number of voxels.

| Classification | Classical  NHST | *γ(Dice_max_)* | *γ = 0.1%* | *γ = 0.2*  *prior SD_θ_* | *γ = 0.5*  *prior SD_θ_* | *γ = 0.8*  *prior SD_θ_* | *γ = 1*  *prior SD_θ_* |
| --- | --- | --- | --- | --- | --- | --- | --- |
| **Switch > baseline. The ‘ROPE-only’ decision rule. *1 prior SD_θ_ = 0.133%*** | | | | | | | |
| Activation | 20.9 | 22.2 | 17.8 | 37.3 | 24.2 | 16.9 | 13.6 |
| Deactivation | 0.7 | 1.3 | 0.6 | 5.2 | 1.7 | 0.5 | 0.2 |
| No activation | - | 36.0 | 52.4 | 0.0 | 28.9 | 55.6 | 66.4 |
| Low-confidence | - | 40.4 | 29.2 | 57.5 | 45.1 | 27.0 | 19.8 |
| **Switch > baseline. The ‘HDI+ROPE’ decision rule. *1 prior SD_θ_ = 0.133%*** | | | | | | | |
| Activation | 20.9 | 22.2 | 16.5 | 34.2 | 22.3 | 15.7 | 12.6 |
| Deactivation | 0.7 | 1.2 | 0.5 | 4.1 | 1.3 | 0.4 | 0.1 |
| No activation | - | 24.2 | 48.0 | 0.0 | 23.6 | 51.4 | 63.3 |
| Low-confidence | - | 52.4 | 35.0 | 61.7 | 52.8 | 32.5 | 24.0 |
| **Switch > No switch. The ‘ROPE-only’ decision rule. *1 prior SD_θ_ = 0.030%*** | | | | | | | |
| Activation | 4.7 | 4.4 | 0.0 | 22.1 | 13.5 | 8.5 | 6.3 |
| Deactivation | 0.0 | 0.0 | 0.0 | 0.8 | 0.1 | 0.0 | 0.0 |
| No activation | - | 59.5 | 97.1 | 0.0 | 0.0 | 23.1 | 42.1 |
| Low-confidence | - | 36.1 | 2.9 | 77.1 | 86.4 | 68.5 | 51.7 |
| **Switch > No switch. The ‘HDI+ROPE’ decision rule. *1 prior SD_θ_ = 0.030%*** | | | | | | | |
| Activation | 4.7 | 4.2 | 0.0 | 17.8 | 10.8 | 6.7 | 4.9 |
| Deactivation | 0.0 | 0.0 | 0.0 | 0.3 | 0.0 | 0.0 | 0.0 |
| No activation | - | 41.5 | 96.4 | 0.0 | 0.0 | 14.7 | 33.0 |
| Low-confidence | - | 54.3 | 3.6 | 81.9 | 89.2 | 78.5 | 62.1 |

### Table S11. Effect sizes for the emotional processing task.

| ROI | Size (voxels) | 25%-tile PSC | Median PSC | 75%-tile PSC | 25%-tile Cohen’s d | Median Cohen’s d | 75%-tile Cohen’s d |
| --- | --- | --- | --- | --- | --- | --- | --- |
| **Emotion processing (Emotional faces > Shapes); *1 prior SD_θ_ = 0.135%*** | | | | | | | |
| L Amy | 324 | 0.174 | 0.263 | 0.407 | 0.557 | 0.744 | 0.968 |
| R Amy | 363 | 0.218 | 0.278 | 0.399 | 0.645 | 0.808 | 1.024 |
| L Fus | 1586 | 0.107 | 0.291 | 0.522 | 0.315 | 0.635 | 0.893 |
| R Fus | 1720 | 0.088 | 0.303 | 0.559 | 0.277 | 0.703 | 0.964 |
| V1 | 3124 | 0.137 | 0.226 | 0.339 | 0.415 | 0.598 | 0.785 |
| V2 | 1335 | 0.122 | 0.186 | 0.272 | 0.353 | 0.520 | 0.679 |

### Table S12. Effect sizes for the working memory task.

| ROI | Size (voxels) | 25%-tile PSC | Median PSC | 75%-tile PSC | 25%-tile Cohen’s d | Median Cohen’s d | 75%-tile Cohen’s d |
| --- | --- | --- | --- | --- | --- | --- | --- |
| **Working memory (2-back > baseline), *1 prior SD_θ_ = 0.325%*** | | | | | | | |
| L DLPFC | 2342 | 0.282 | 0.422 | 0.597 | 0.581 | 0.800 | 0.990 |
| R DLPFC | 2539 | 0.250 | 0.388 | 0.541 | 0.560 | 0.767 | 0.985 |
| L SPL | 821 | 0.216 | 0.418 | 0.608 | 0.674 | 1.045 | 1.340 |
| R SPL | 524 | 0.262 | 0.392 | 0.520 | 0.699 | 1.029 | 1.276 |
| L IPL | 876 | 0.202 | 0.334 | 0.470 | 0.769 | 1.004 | 1.251 |
| R IPL | 1023 | 0.211 | 0.300 | 0.426 | 0.752 | 0.990 | 1.214 |
| dACC | 1184 | 0.295 | 0.469 | 0.751 | 0.589 | 0.867 | 1.235 |
| V1 | 3165 | 0.287 | 0.536 | 0.879 | 0.590 | 0.857 | 1.104 |
| V2 | 1356 | 0.199 | 0.375 | 0.861 | 0.361 | 0.621 | 0.972 |
| **Working memory (2-back > 0-back), *1 prior SD_θ_ = 0.089%*** | | | | | | | |
| L DLPFC | 2342 | 0.090 | 0.133 | 0.189 | 0.279 | 0.412 | 0.582 |
| R DLPFC | 2539 | 0.075 | 0.137 | 0.207 | 0.263 | 0.464 | 0.638 |
| L SPL | 821 | 0.029 | 0.073 | 0.144 | 0.156 | 0.325 | 0.534 |
| R SPL | 524 | 0.021 | 0.079 | 0.144 | 0.105 | 0.346 | 0.587 |
| L IPL | 876 | 0.096 | 0.145 | 0.195 | 0.494 | 0.664 | 0.769 |
| R IPL | 1023 | 0.128 | 0.176 | 0.223 | 0.593 | 0.762 | 0.887 |
| dACC | 1184 | 0.099 | 0.174 | 0.260 | 0.304 | 0.465 | 0.609 |
| V1 | 3165 | -0.043 | -0.009 | 0.019 | -0.150 | -0.034 | 0.069 |
| V2 | 1356 | -0.037 | -0.011 | 0.016 | -0.134 | -0.041 | 0.054 |

### Table S13. Effect sizes for the language task.

| ROI | Size (voxels) | 25%-tile PSC | Median PSC | 75%-tile PSC | 25%-tile Cohen’s d | Median Cohen’s d | 75%-tile Cohen’s d |
| --- | --- | --- | --- | --- | --- | --- | --- |
| **Language (Story > Math), *1 prior SD_θ_ = 0.255%*** | | | | | | | |
| L IFG pt  (BA 45) | 785 | 0.011 | 0.269 | 0.525 | 0.027 | 0.476 | 0.876 |
| L IFG po  (BA 44) | 808 | -0.318 | -0.104 | 0.063 | -0.593 | -0.246 | 0.136 |
| L STG  (Wernicke) | 1316 | 0.135 | 0.292 | 0.542 | 0.450 | 0.785 | 1.014 |
| L Heschl | 408 | 0.151 | 0.235 | 0.309 | 0.486 | 0.693 | 0.848 |
| R Heschl | 378 | 0.103 | 0.184 | 0.272 | 0.367 | 0.626 | 0.856 |

### Table S14. Effect sizes for the motor task.

| ROI | Size (voxels) | 25%-tile PSC | Median PSC | 75%-tile PSC | 25%-tile Cohen’s d | Median Cohen’s d | 75%-tile Cohen’s d |
| --- | --- | --- | --- | --- | --- | --- | --- |
| **Motor (Left finger > baseline), *1 prior SD_θ_ = 0.149%*** | | | | | | | |
| L Precent | 3198 | -0.036 | 0.057 | 0.159 | -0.097 | 0.165 | 0.426 |
| R Precent | 2705 | 0.081 | 0.194 | 0.338 | 0.239 | 0.504 | 0.783 |
| L Postcent | 1882 | -0.145 | -0.028 | 0.151 | -0.438 | -0.090 | 0.416 |
| R Postcent | 1318 | 0.228 | 0.330 | 0.504 | 0.606 | 0.840 | 1.078 |
| L Put | 672 | 0.097 | 0.152 | 0.207 | 0.176 | 0.293 | 0.384 |
| R Put | 659 | 0.079 | 0.197 | 0.284 | 0.134 | 0.356 | 0.535 |
| L Cerebell | 1139 | 0.200 | 0.352 | 0.568 | 0.391 | 0.661 | 1.051 |
| R Cerebell | 1604 | 0.031 | 0.110 | 0.221 | 0.060 | 0.227 | 0.479 |
| SMA | 1639 | 0.100 | 0.234 | 0.401 | 0.237 | 0.466 | 0.703 |
| V1 | 3172 | -0.084 | -0.030 | 0.022 | -0.154 | -0.059 | 0.049 |
| V2 | 1366 | -0.105 | -0.046 | 0.044 | -0.208 | -0.098 | 0.084 |
| **Motor (Right finger > baseline), *1 prior SD_θ_ = 0.171%*** | | | | | | | |
| L Precent | 3198 | 0.116 | 0.239 | 0.375 | 0.294 | 0.545 | 0.816 |
| R Precent | 2705 | 0.002 | 0.111 | 0.235 | 0.006 | 0.278 | 0.535 |
| L Postcent | 1882 | 0.245 | 0.362 | 0.553 | 0.630 | 0.837 | 1.072 |
| R Postcent | 1318 | -0.129 | -0.050 | 0.079 | -0.383 | -0.144 | 0.209 |
| L Put | 672 | 0.194 | 0.290 | 0.370 | 0.305 | 0.454 | 0.565 |
| R Put | 659 | 0.069 | 0.122 | 0.184 | 0.104 | 0.189 | 0.273 |
| L Cerebell | 1139 | 0.104 | 0.207 | 0.329 | 0.205 | 0.400 | 0.622 |
| R Cerebell | 1604 | 0.224 | 0.401 | 0.573 | 0.411 | 0.671 | 0.904 |
| SMA | 1639 | 0.175 | 0.353 | 0.559 | 0.373 | 0.608 | 0.827 |
| V1 | 3172 | -0.093 | -0.027 | 0.032 | -0.172 | -0.055 | 0.067 |
| V2 | 1366 | -0.088 | -0.026 | 0.087 | -0.161 | -0.050 | 0.172 |
| **Motor (Tongue > baseline), *1 prior SD_θ_ = 0.268%*** | | | | | | | |
| L Precent | 3198 | 0.076 | 0.288 | 0.554 | 0.152 | 0.551 | 0.956 |
| R Precent | 2705 | 0.013 | 0.234 | 0.520 | 0.024 | 0.435 | 0.860 |
| L Postcent | 1882 | -0.041 | 0.163 | 0.586 | -0.104 | 0.358 | 1.009 |
| R Postcent | 1318 | -0.102 | 0.028 | 0.348 | -0.253 | 0.061 | 0.652 |
| L Put | 672 | 0.032 | 0.166 | 0.245 | 0.028 | 0.180 | 0.269 |
| R Put | 659 | -0.182 | -0.008 | 0.104 | -0.187 | -0.008 | 0.106 |
| L Cerebell | 1139 | 0.075 | 0.239 | 0.528 | 0.084 | 0.298 | 0.682 |
| R Cerebell | 1604 | 0.056 | 0.218 | 0.455 | 0.054 | 0.261 | 0.599 |
| SMA | 1639 | 0.079 | 0.292 | 0.579 | 0.121 | 0.401 | 0.697 |
| V1 | 3172 | -0.156 | -0.074 | -0.014 | -0.240 | -0.128 | -0.025 |
| V2 | 1366 | -0.169 | -0.085 | 0.024 | -0.272 | -0.142 | 0.036 |

### Table S15. Effect sizes for the gambling task.

| ROI | Size (voxels) | | 25%-tile PSC | Median PSC | 75%-tile PSC | 25%-tile Cohen’s d | Median Cohen’s d | 75%-tile Cohen’s d |
| --- | --- | --- | --- | --- | --- | --- | --- | --- |
| **Gambling (Reward > baseline), *1 prior SD_θ_ = 0.254%*** | | | | | | | | |
| L NAc | 55 | | 0.002 | 0.029 | 0.069 | 0.004 | 0.063 | 0.150 |
| R NAc | 64 | | -0.014 | 0.035 | 0.083 | -0.033 | 0.079 | 0.183 |
| L Amy | 308 | | -0.137 | -0.096 | -0.050 | -0.337 | -0.240 | -0.132 |
| R Amy | 348 | | -0.127 | -0.101 | -0.062 | -0.320 | -0.257 | -0.147 |
| L Ins | 889 | | 0.021 | 0.114 | 0.276 | 0.059 | 0.320 | 0.763 |
| R Ins | 853 | | 0.000 | 0.090 | 0.248 | 0.001 | 0.272 | 0.649 |
| V1 | 3164 | | 0.430 | 0.716 | 1.017 | 0.842 | 1.204 | 1.452 |
| V2 | 1369 | | 0.456 | 0.654 | 0.878 | 0.830 | 1.054 | 1.317 |
| **Gambling (Loss > baseline), *1 prior SD_θ_ = 0.249%*** | | | | | | | | |
| L NAc | 55 | -0.190 | | -0.152 | -0.111 | -0.428 | -0.373 | -0.245 |
| R NAc | 64 | -0.131 | | -0.103 | -0.087 | -0.286 | -0.229 | -0.192 |
| L Amy | 308 | -0.208 | | -0.165 | -0.112 | -0.463 | -0.389 | -0.287 |
| R Amy | 348 | -0.178 | | -0.124 | -0.072 | -0.402 | -0.301 | -0.187 |
| L Ins | 889 | -0.008 | | 0.083 | 0.217 | -0.025 | 0.233 | 0.561 |
| R Ins | 853 | -0.028 | | 0.057 | 0.214 | -0.079 | 0.158 | 0.566 |
| V1 | 3164 | 0.389 | | 0.656 | 0.924 | 0.769 | 1.117 | 1.369 |
| V2 | 1369 | 0.402 | | 0.559 | 0.787 | 0.739 | 0.943 | 1.208 |
| **Gambling (Reward > Loss), *1 prior SD_θ_ = 0.032%*** | | | | | | | | |
| L NAc | 55 | | 0.031 | 0.043 | 0.048 | 0.103 | 0.136 | 0.154 |
| R NAc | 64 | | 0.027 | 0.034 | 0.042 | 0.093 | 0.110 | 0.136 |
| L Amy | 308 | | 0.009 | 0.018 | 0.025 | 0.034 | 0.060 | 0.086 |
| R Amy | 348 | | -0.003 | 0.008 | 0.018 | -0.011 | 0.027 | 0.062 |
| L Ins | 889 | | 0.001 | 0.014 | 0.030 | 0.003 | 0.054 | 0.120 |
| R Ins | 853 | | 0.002 | 0.012 | 0.021 | 0.010 | 0.049 | 0.083 |
| V1 | 3164 | | 0.012 | 0.027 | 0.047 | 0.053 | 0.115 | 0.200 |
| V2 | 1369 | | 0.007 | 0.027 | 0.067 | 0.028 | 0.115 | 0.281 |

### Table S16. Effect sizes for the social cognition task.

| ROI | Size (voxels) | | 25%-tile PSC | Median PSC | 75%-tile PSC | 25%-tile Cohen’s d | Median Cohen’s d | 75%-tile Cohen’s d |
| --- | --- | --- | --- | --- | --- | --- | --- | --- |
| **Social cognition (Social interaction > baseline), *1 prior SD_θ_ = 0.325%*** | | | | | | | | |
| L IPL | 1158 | | 0.142 | 0.288 | 0.491 | 0.486 | 0.796 | 1.049 |
| R IPL | 1421 | | 0.166 | 0.435 | 0.653 | 0.486 | 0.916 | 1.222 |
| L STG | 477 | | 0.096 | 0.190 | 0.284 | 0.295 | 0.487 | 0.605 |
| R STG | 548 | | 0.210 | 0.348 | 0.477 | 0.456 | 0.679 | 0.880 |
| L MTG | 872 | | 0.097 | 0.303 | 0.537 | 0.294 | 0.683 | 1.107 |
| R MTG | 937 | | 0.208 | 0.465 | 0.677 | 0.518 | 0.921 | 1.244 |
| V1 | 3163 | | 0.077 | 0.253 | 0.600 | 0.168 | 0.437 | 0.900 |
| V2 | 1352 | | -0.118 | 0.018 | 0.321 | -0.189 | 0.029 | 0.447 |
| **Social cognition (Social interaction > Random interaction), *1 prior SD_θ_ = 0.104%*** | | | | | | | | |
| L IPL | 1158 | 0.059 | | 0.142 | 0.209 | 0.217 | 0.489 | 0.665 |
| R IPL | 1421 | -0.009 | | 0.137 | 0.233 | -0.037 | 0.476 | 0.697 |
| L STG | 477 | 0.122 | | 0.184 | 0.224 | 0.398 | 0.529 | 0.626 |
| R STG | 548 | 0.159 | | 0.252 | 0.325 | 0.440 | 0.615 | 0.744 |
| L MTG | 872 | 0.109 | | 0.169 | 0.218 | 0.331 | 0.560 | 0.720 |
| R MTG | 937 | 0.127 | | 0.212 | 0.274 | 0.409 | 0.660 | 0.804 |
| V1 | 3163 | -0.035 | | 0.013 | 0.051 | -0.090 | 0.032 | 0.123 |
| V2 | 1352 | -0.118 | | -0.030 | 0.030 | -0.300 | -0.076 | 0.070 |

### Table S17. Effect sizes for the relational processing task.

| ROI | Size (voxels) | 25%-tile PSC | Median PSC | 75%-tile PSC | 25%-tile Cohen’s d | Median Cohen’s d | 75%-tile Cohen’s d |
| --- | --- | --- | --- | --- | --- | --- | --- |
| **Relational processing (Relational > baseline), *1 prior SD_θ_ = 0.390%*** | | | | | | | |
| L DLPFC | 646 | 0.463 | 0.681 | 0.980 | 0.859 | 1.063 | 1.205 |
| R DLPFC | 334 | 0.363 | 0.513 | 0.673 | 0.757 | 0.924 | 1.130 |
| L SPL | 168 | 0.721 | 0.842 | 0.943 | 1.424 | 1.536 | 1.658 |
| R SPL | 37 | 0.574 | 0.661 | 0.728 | 1.239 | 1.359 | 1.434 |
| L AI/FO | 360 | 0.228 | 0.410 | 0.643 | 0.488 | 0.813 | 1.151 |
| R AI/FO | 198 | 0.289 | 0.496 | 0.700 | 0.673 | 1.031 | 1.284 |
| preSMA/dACC | 536 | 0.562 | 0.851 | 1.158 | 0.887 | 1.123 | 1.351 |
| V1 | 3131 | 0.742 | 1.152 | 1.582 | 1.065 | 1.384 | 1.628 |
| V2 | 1341 | 0.707 | 0.987 | 1.340 | 0.961 | 1.157 | 1.488 |
| **Relational processing (Relational > Match), *1 prior SD_θ_ = 0.051%*** | | | | | | | |
| L DLPFC | 646 | 0.043 | 0.062 | 0.077 | 0.153 | 0.219 | 0.297 |
| R DLPFC | 334 | 0.024 | 0.058 | 0.084 | 0.114 | 0.234 | 0.315 |
| L SPL | 168 | -0.021 | 0.003 | 0.026 | -0.088 | 0.012 | 0.102 |
| R SPL | 37 | 0.026 | 0.059 | 0.084 | 0.122 | 0.269 | 0.414 |
| L AI/FO | 360 | 0.012 | 0.038 | 0.072 | 0.037 | 0.131 | 0.233 |
| R AI/FO | 198 | 0.017 | 0.044 | 0.069 | 0.057 | 0.141 | 0.216 |
| preSMA/dACC | 536 | 0.051 | 0.081 | 0.109 | 0.154 | 0.233 | 0.297 |
| V1 | 3131 | -0.040 | -0.009 | 0.055 | -0.142 | -0.035 | 0.184 |
| V2 | 1341 | -0.051 | 0.006 | 0.063 | -0.186 | 0.021 | 0.207 |

### Table S18. Effect sizes for the stop-signal task.

| ROI | Size (voxels) | 25%-tile PSC | Median PSC | 75%-tile PSC | 25%-tile Cohen’s d | Median Cohen’s d | 75%-tile Cohen’s d |
| --- | --- | --- | --- | --- | --- | --- | --- |
| **Stop-signal task (Correct Stop > baseline), *1 prior SD_θ_ = 0.069%*** | | | | | | | |
| L IFG/FO | 116 | -0.066 | 0.093 | 0.173 | -0.266 | 0.408 | 0.587 |
| R IFG/FO | 272 | 0.047 | 0.095 | 0.149 | 0.220 | 0.407 | 0.575 |
| L DLPFC | 143 | -0.049 | 0.030 | 0.056 | -0.206 | 0.147 | 0.258 |
| R DLPFC | 349 | 0.024 | 0.057 | 0.102 | 0.093 | 0.241 | 0.430 |
| preSMA/dACC | 308 | 0.051 | 0.089 | 0.126 | 0.246 | 0.365 | 0.475 |
| L Heschl | 110 | 0.073 | 0.108 | 0.144 | 0.330 | 0.484 | 0.605 |
| R Heschl | 110 | 0.053 | 0.085 | 0.144 | 0.251 | 0.377 | 0.520 |
| V1 | 796 | -0.105 | -0.028 | 0.010 | -0.325 | -0.100 | 0.045 |
| V2 | 417 | -0.131 | -0.052 | -0.009 | -0.397 | -0.177 | -0.033 |
| **Stop-signal task (Correct Stop > Go), *1 prior SD_θ_ = 0.064%*** | | | | | | | |
| L IFG/FO | 116 | 0.020 | 0.071 | 0.182 | 0.112 | 0.403 | 0.812 |
| R IFG/FO | 272 | 0.080 | 0.120 | 0.162 | 0.497 | 0.626 | 0.767 |
| L DLPFC | 143 | 0.024 | 0.038 | 0.051 | 0.151 | 0.219 | 0.297 |
| R DLPFC | 349 | 0.065 | 0.092 | 0.118 | 0.420 | 0.524 | 0.617 |
| preSMA/dACC | 308 | 0.058 | 0.090 | 0.132 | 0.377 | 0.515 | 0.681 |
| L Heschl | 110 | 0.068 | 0.111 | 0.142 | 0.386 | 0.662 | 0.791 |
| R Heschl | 110 | 0.065 | 0.107 | 0.164 | 0.353 | 0.614 | 0.736 |
| V1 | 796 | 0.028 | 0.069 | 0.100 | 0.151 | 0.322 | 0.425 |
| V2 | 417 | 0.014 | 0.043 | 0.076 | 0.069 | 0.201 | 0.342 |

### Table S19. Effect sizes for the task-switching task.

| ROI | Size (voxels) | 25%-tile PSC | Median PSC | 75%-tile PSC | 25%-tile Cohen’s d | Median Cohen’s d | 75%-tile Cohen’s d |
| --- | --- | --- | --- | --- | --- | --- | --- |
| **Task-switching (Switch > baseline), *1 prior SD_θ_ = 0.133%*** | | | | | | | |
| L DLPFC | 157 | 0.148 | 0.206 | 0.277 | 0.471 | 0.599 | 0.716 |
| R DLPFC | 209 | 0.066 | 0.095 | 0.131 | 0.234 | 0.321 | 0.403 |
| L SPL | 151 | 0.194 | 0.323 | 0.430 | 0.688 | 0.956 | 1.073 |
| R SPL | 63 | 0.148 | 0.218 | 0.272 | 0.530 | 0.669 | 0.756 |
| L LOC | 258 | 0.152 | 0.292 | 0.407 | 0.525 | 0.780 | 0.945 |
| R LOC | 162 | 0.149 | 0.218 | 0.292 | 0.516 | 0.678 | 0.812 |
| preSMA/dACC | 340 | 0.106 | 0.152 | 0.199 | 0.383 | 0.497 | 0.595 |
| L AI/FO | 130 | 0.053 | 0.105 | 0.165 | 0.251 | 0.376 | 0.557 |
| R AI/FO | 94 | 0.077 | 0.124 | 0.162 | 0.286 | 0.442 | 0.543 |
| V1 | 791 | 0.053 | 0.121 | 0.310 | 0.179 | 0.349 | 0.614 |
| V2 | 411 | 0.084 | 0.193 | 0.429 | 0.253 | 0.455 | 0.899 |
| **Task switching (Switch > No switch), *1 prior SD_θ_ = 0.030%*** | | | | | | | |
| L DLPFC | 157 | 0.056 | 0.074 | 0.088 | 0.372 | 0.446 | 0.486 |
| R DLPFC | 209 | 0.028 | 0.039 | 0.053 | 0.216 | 0.284 | 0.345 |
| L SPL | 151 | 0.047 | 0.079 | 0.104 | 0.377 | 0.523 | 0.624 |
| R SPL | 63 | 0.029 | 0.046 | 0.061 | 0.225 | 0.330 | 0.386 |
| L LOC | 258 | 0.054 | 0.081 | 0.102 | 0.363 | 0.469 | 0.546 |
| R LOC | 162 | 0.031 | 0.044 | 0.061 | 0.232 | 0.333 | 0.409 |
| preSMA/dACC | 340 | 0.043 | 0.056 | 0.075 | 0.313 | 0.400 | 0.453 |
| L AI/FO | 130 | 0.036 | 0.056 | 0.076 | 0.306 | 0.418 | 0.545 |
| R AI/FO | 94 | 0.037 | 0.050 | 0.066 | 0.284 | 0.383 | 0.492 |
| V1 | 791 | 0.014 | 0.024 | 0.035 | 0.091 | 0.157 | 0.216 |
| V2 | 411 | 0.008 | 0.017 | 0.026 | 0.053 | 0.109 | 0.170 |
